# Regulatory mechanisms in multiple vascular diseases locus *LRP1* involve repression by SNAIL and extracellular matrix remodeling

**DOI:** 10.1101/2023.05.09.539992

**Authors:** Lu Liu, Joséphine Henry, Yingwei Liu, Charlène Jouve, Adrien Georges, Nabila Bouatia-Naji

**Affiliations:** Université Paris Cité, Inserm, PARCC, F-75015 Paris, France

## Abstract

**Background:** Genome-wide association studies (GWAS) implicate common genetic variations in the low-density lipoprotein receptor-related protein 1 locus *(LRP1)* in risk for multiple vascular diseases and traits. However, the underlying biological mechanisms are unknown.

**Methods:** Fine mapping analyses included Bayesian colocalization to identify the most likely causal variant. Human induced pluripotent stem cells (iPSC) were genome-edited using CRISPR-Cas9 to delete or modify candidate enhancer regions, and generate *LRP1* knockout cell lines (KO). Cells were differentiated into smooth muscle cells (SMCs) through a mesodermal lineage. Transcription regulation was assessed using luciferase reporter assay, transcription factor knockdown and chromatin immunoprecipitation. Phenotype changes in cells were conducted using cellular assays, bulk RNA-sequencing and mass spectrometry.

**Results:** Multi-trait co-localization analyses pointed at rs11172113 as the most likely causal variant in *LRP1* for fibromuscular dysplasia, migraine, pulse pressure and pulmonary function trait. We found rs11172113-T allele to associate with higher *LRP1* expression. Genomic deletion in iPSC-derived SMCs supported rs11172113 to locate in an enhancer region regulating *LRP1* expression. We found transcription factors MECP2 and SNAIL to repress *LRP1* expression through an allele-specific mechanism, involving SNAIL interaction with disease risk allele. *LRP1* KO decreased iPSC-derived SMCs proliferation and migration. Differentially expressed genes were enriched for collagen-containing extracellular matrix, connective tissue and lung development. *LRP1* KO showed potentiated canonical TGF-β signaling through enhanced phosphorylation of SMAD2/3. Analyses of protein content of decellularized extracts indicated partial extracellular matrix (ECM) remodeling involving enhanced secretion of CYR61, a known LRP1 ligand involved in vascular integrity and TIMP3, implicated in extracellular matrix maintenance and also known to interact with LRP1.

**Conclusions:** Our findings support allele specific *LRP1* gene repression by the endothelial-to-mesenchymal transition regulator SNAIL. We propose decreased *LRP1* expression in SMCs to remodel the ECM enhanced by TGF- β as a potential mechanism of this pleiotropic locus for vascular diseases.

**Graphical abstract:** 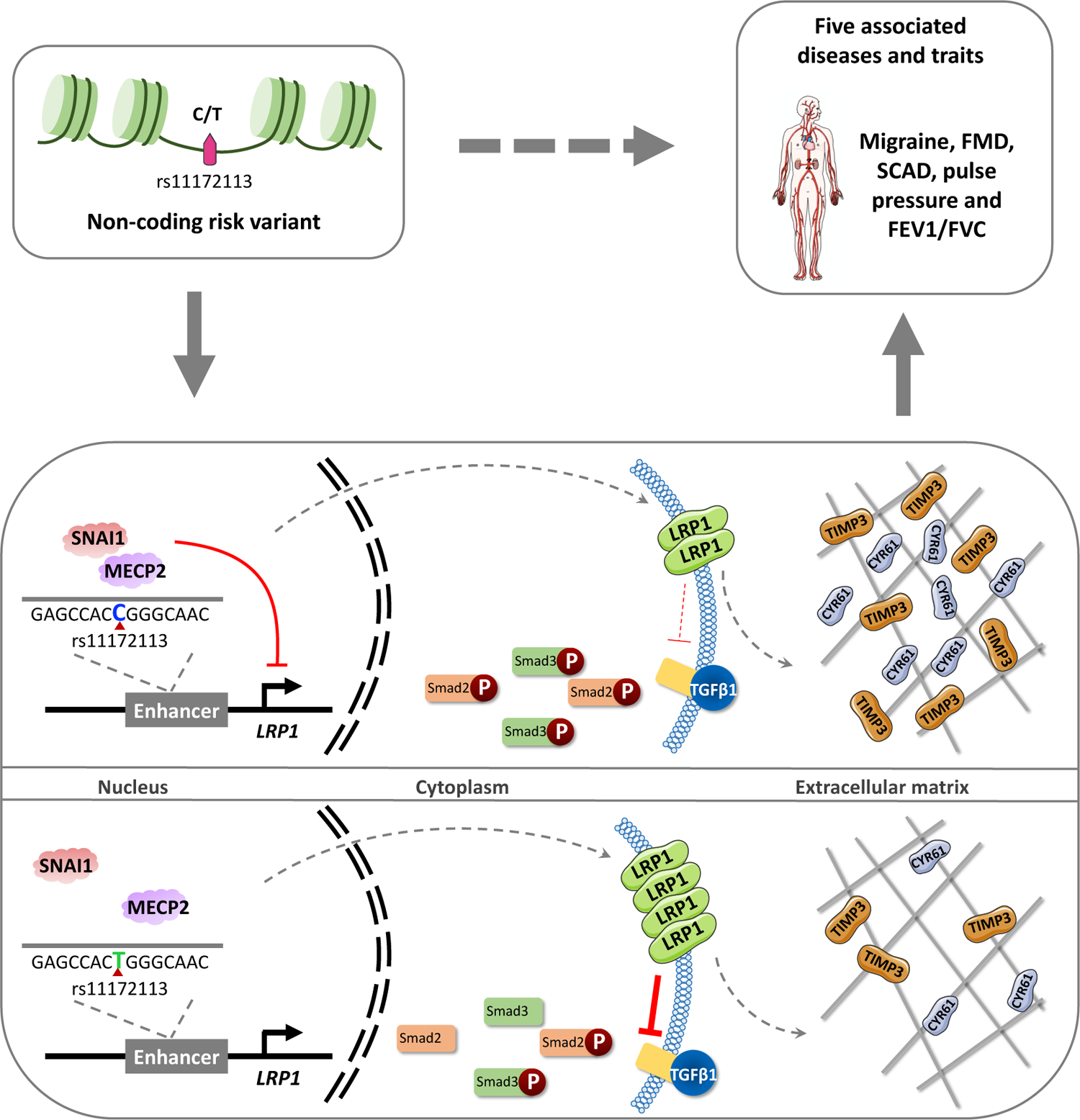

## Introduction

Genome-wide association studies (GWAS) have established thousands of genomic locations in link with cardiovascular diseases.^1–4^ GWAS generally produces complex association signals that are difficult to transcribe into clear biological mechanisms. Identified risk loci in GWAS are often highly pleiotropic involving association signals related to several diseases and traits that may potentially share unknown molecular and cellular mechanisms.^5^ Causal variants at GWAS loci are difficult to determine due to high linkage disequilibrium between associated variants that results in the redundancy of statistical signals in most loci. In addition, associated variants are mostly non-coding,^6, 7^ and likely to be involved in tissue or cell type specific gene expression regulation^8^. The identification of causal variations and their biological significance is particularly challenging in the case of inaccessible tissues such as arteries.

Several GWAS have reported robust association signals on chromosome 12, in the first intron of the low-density lipoprotein receptor-related protein 1 gene (*LRP1*). First reported in migraine^9^, *LRP1* was also described as a risk locus for sporadic thoracic aortic dissection^10^, and more recently for fibromuscular dysplasia (FMD), an arteriopathy associated with resistant hypertension and stroke, and with an underdiagnosed form of ischemic heart disease named spontaneous coronary artery dissection (SCAD)^11, 12^. *LRP1* locus is also associated with pulse pressure (PP)^4^, a cardiovascular risk factor indicative of arterial stiffness^13^, and forced expiratory volume over vital capacity (FEV1/FVC)^14^, a surrogate trait of obstructive lung disease^15^. The molecular and cellular mechanisms underlying these genetic associations with this wide range of vascular diseases and traits are still to be determined.

LRP1 is an endocytic transmembrane protein member of the low-density lipoprotein receptor family that interacts with numerous biologically diverse ligands^16^ and was extensively studied in the context of several human diseases. LRP1 is involved in many cellular processes through its capacity to internalize a wide variety of extracellular proteins. *LRP1* regulates the composition of the extracellular matrix and interacts with key signaling pathways such as transforming growth factor beta (TGF-β) or platelet-derived growth factor (PDGF) pathways^17^. LRP1 is highly expressed in smooth muscle cells (SMCs) where it was demonstrated to maintain endocytosis and vascular integrity^18^. LRP1 regulates SMCs proliferation, migration^19^, extracellular matrix deposition,^20^ in addition to controlling calcium efflux to maintain arterial contractility^21^. Independent studies in mice and human cells suggested a role for SMC-expressed *LRP1* in atherosclerosis^19, 22^ and pulmonary hypertension^23^, with conflicting reports on the cellular outcome of LRP1 deficiency^19, 22, 23^.

In this integrative study, we report plausible regulatory mechanisms for the associations observed at the *LRP1* locus. We leveraged multiple trait associations at this highly pleiotropic locus to formally identify rs11172113 as the most likely causal variant for reported associations. We assessed the biological function of this variant using genetically modified SMCs derived from human induced pluripotent stem cells (iPSCs) through a mesoderm lineage. We confirmed rs11172113 to belong to an enhancer region intronic to *LRP1*, and provide data supporting transcriptional repressors, SNAIL and MECP2, to regulate *LRP1* expression in an allele-specific manner. We showed the binding of SNAIL to this enhancer was impaired by risk allele for diseases. Finally, we found that cellular knockout of *LRP1* in iPSC-derived SMCs led to hyperactivation of TGF-β signaling and hypersecretion of TIMP3 and CYR61, two LRP1 ligands involved in vascular integrity.

## Results

### *LRP1* is a pleiotropic locus involved in the risk of multiple arterial diseases

*LRP1* locus was previously reported to associate with migraine^24^ (OR=1.11, *P*=1×10^-90^), FMD^11^ (OR=1.33, *P*=2×10^-10^), SCAD^12^ (OR=1.51, *P*=3×10^-13^), sporadic thoracic aortic dissection^10^ (OR=1.21, *P*=3×10^-8^), and suggestively to cervical artery dissection (CeAD)^25^ (OR=1.22, *P*=3×10^-7^), as summarized in **Figure 1A**. We applied conditional regression analyses using GCTA COJO method,^26^ and confirmed a single genetic signal that involves rs11172113 as the lead variant in FMD and migraine signals, for which we had access to full summary statistics (**Figure 1B, Figure S1**). Multi-trait Bayesian colocalization analysis^27^ of association signals for migraine, FMD, pulse pressure and FEV1/FVC indicated high posterior probabilities for colocalization with a single shared causal variant (PP_4 traits_=99.3%, **Figure 1C**), supporting rs11172113 to likely explain the observed association signals (**Figure 1C**). In the same line, colocalization analyses between GWAS and expression quantitative trait loci (eQTLs) signals in aorta, tibial and coronary arteries from GTEx also supported rs111712113 as the likely causal variant in this locus in these tissues, with *LRP1* as the most likely and only target gene (PP.H4.abf >80%, **Figure 1D, Figure S2**). Of note, no eQTL association signal was observed for rs11172113 in pulmonary tissues, although a suggestive eQTL signal, colocalizing with arterial eQTL, is found in fibroblasts (**Figure S2**). Altogether, our results support rs11172113 as the most likely causal variant and candidate functional variant potentially involved in *LRP1* expression in arteries.

**Figure 1.**
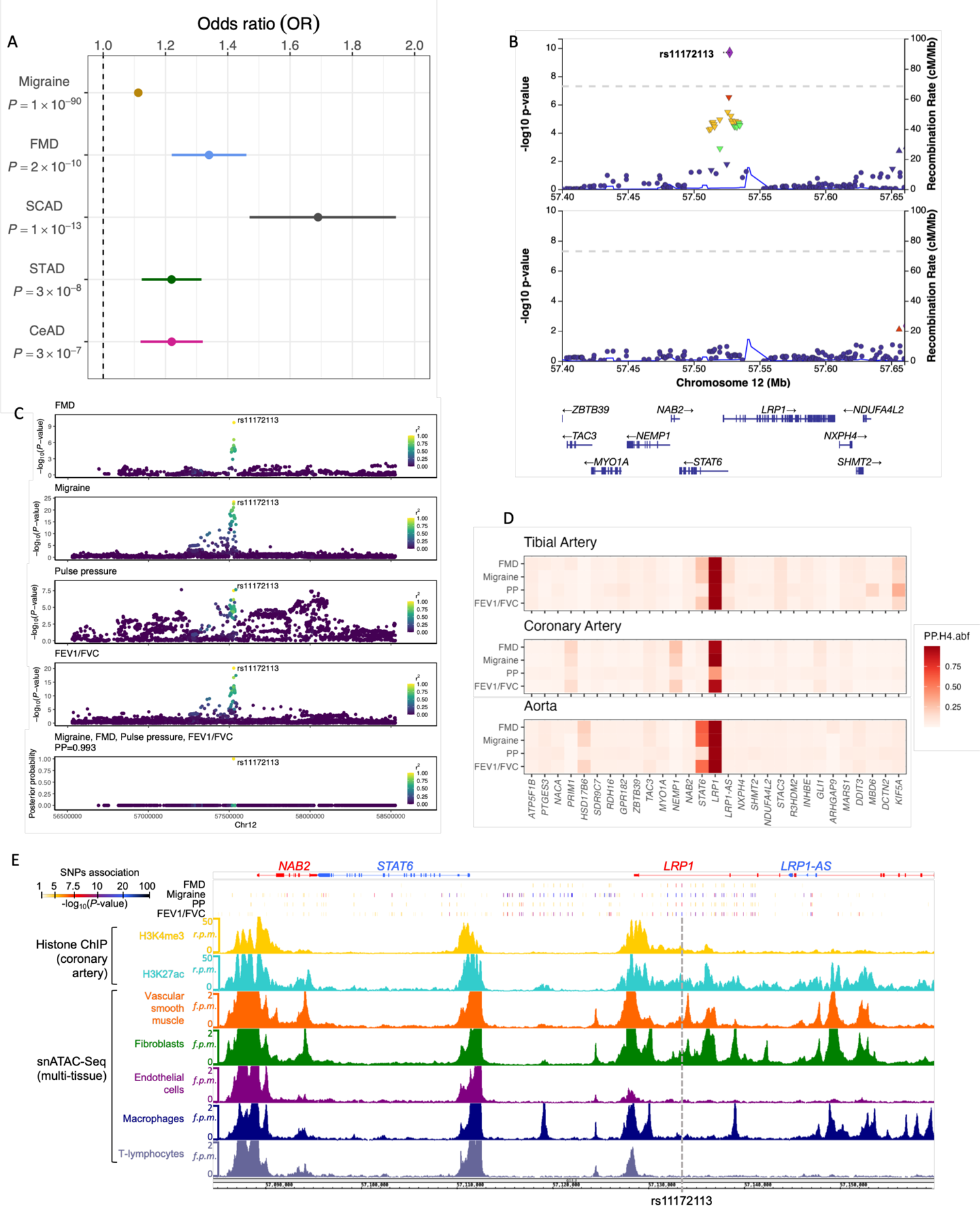
Genetic associations and functional annotations at the *LRP1* locus. **A**: Forest plot representing odds ratios (OR) estimations and p-values (*P*) of the associations between rs11172113 (T allele at risk) and migraine, fibromuscular dysplasia (FMD), spontaneous coronary artery dissection (SCAD), sporadic thoracic aorta dissection (STAD) and cervical artery dissection (CeAD). **B**: Representative locusZoom plots of FMD association signal before (top) and after (bottom) conditioning on rs11172113 association using GCTA COJO method^26^. **C**: Multitrait colocalization analysis performed using HyPrColoc method ^27^. Genetic association signals for Migraine, FMD and FEV1/FVC are represented in a 2Mb region centered on rs11172113. Association P-value (log10 scale) of each SNP is represented on y-axis, while dot color represents r^2^ of linkage disequilibrium with rs11172113 (European population of 1000G reference panel). Lower panel represents the relative posterior probability for each SNP to be causal, while the total posterior probability for the traits to colocalize at the locus is given over the graph (PP=0.993) **D**: Heatmap representing colocalizations of genetic association signals for 4 traits or diseases, and eQTL association signals in artery tissues. All genes overlapping a 1Mb window centered on rs11172113 were tested for colocalization. Tile color represents the H4 coefficient of approximate Bayes factor colocalization (PP.H4.abf, 0 to 1). **E**: Genome browser visualization of histone chromatin-immunoprecipitation (Histone-ChIP) and single nuclei snATAC-Seq read densities in reads per million (r.p.m) in the regions surrounding rs11172113. Dashed grey line highlights rs11172113 exact position.

### rs11172113 is located in an enhancer region active in iPSC-derived SMCs

We have previously shown that rs11172113 is located within an open chromatin region in primary arterial smooth muscle cells and dermal fibroblasts^11^. Using open chromatin maps generated by single nuclei ATAC-seq from multiple tissues^28^, we confirmed rs11172113 to overlap with open chromatin regions specifically in vascular SMCs and fibroblasts, but not in endothelial cells, macrophages or T-lymphocytes (**Figure 1E**). To assay the enhancer activity of this genomic region, we cloned a 1 kb genomic region centered on rs11172113 (T or C allele) in a reporter plasmid (**Figure 2A**). Enhancer dual-luciferase reporter assays indicated that this sequence produced significantly higher luciferase activity compared to a control sequence, both in rat (**Figure 2B**) and iPSC-derived SMCs (**Figure 2C**). However, we observed no significant differences in luciferase activities between T and C alleles constructs (**Figure 2B-C**).

**Figure 2.**
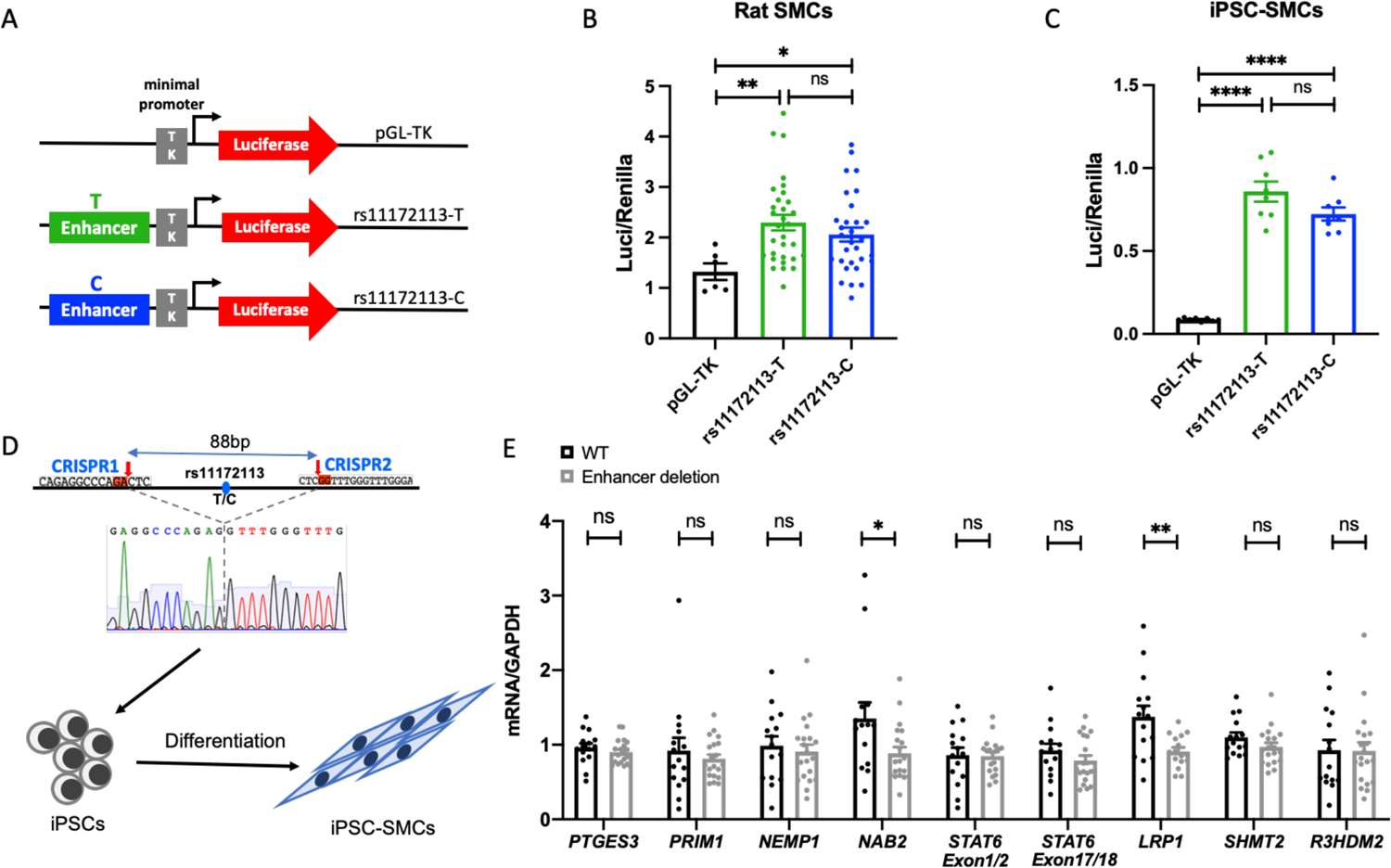
Enhancer capacity of genomic region harboring rs11172113. **A**: Illustration of luciferase plasmids constructs analyzed including rs11172113-centered region for each allele (T/C) orthogonally inserted into the pGL-TK luciferase vector with minimal thymidine kinase promoter (TK). **B-C**: Dual luciferase assay for enhancer activity of empty vectors (gray), and plasmids with 1kb region centered on rs11172113 with the T (green) or C allele (blue) in rat (**B**) or iPSC-SMCs (**C**). Unadjusted p-values of comparison between sample groups using Student’s t-test are indicated: mean±SEM, ****p<10^-4^, ns: not significant. **D**: Illustration of the 100bp enhancer region centered on rs11172113 that was deleted in iPSCs, which were subsequently differentiated into SMCs (iPSC-SMCs). **E**: Relative expression of genes in the vicinity of LRP1 locus in iPSC-SMCs with or without rs11172113-associated enhancer. Four WT and five enhancer deletion clones were analyzed in four independent experiments, which were grouped. Bar plot represent mean±SEM. Unadjusted t-test: *p<0.05, **p<0.01, ns: not significant.

To further confirm the enhancer activity of rs11172113 surrounding region, we generated iPSCs where we deleted using the CRISPR-Cas9 system an 88 bp sequence centered on rs11172113 (**Figure 2D**). We obtained five single clones with sequence deletions, which we compared to four single clones without deletion generated using Cas9 overexpression in absence of gRNA (**Table S1**). Compared to their parental cell line, all clones presented similar expressions of pluripotency markers (**Figure S3**). We differentiated all clones into SMCs throughout the mesodermal lineage as previously described^29^. We found a significant decrease in *LRP1* expression in SMCs encompassing the deletion, compared to wild type SMCs, supporting an enhancer effect of this region on the expression of *LRP1* (**Figure 2E**). We also examined the expression of genes mapping in the vicinity of *LRP1* (within ±0.3 Mb and reported to be expressed in arteries). In addition to *LRP1*, we found a significant decrease in the expression of the *NGFI-A binding protein 2* gene (*NAB2*), located 40kb upstream of rs11172113 (**Figure 2E**). rs11172113 is reported in GTEx as a splicing QTL in tibial artery with allele T associated to a lower inclusion of canonical exon 1-exon 2 junction within the transcript for signal transducer and activator of transcription 6 gene (*STAT6*) (**Figure S4**). However, we did not observe changes in the expression of *STAT6* when we targeted exon 1-exon 2 and exon 17 - exon 18 junctions (**Figure 2E**). This result does not support rs11172113 variation as the causal variant for this sQTL described in GTEx.

### Transcription factors MECP2 and SNAIL are repressors of *LRP1* expression

We used PERFECTOS-APE^30^ to assess potential changes in the binding capacity or recruitment of transcriptional factors (TFs) according to rs11172113 alleles. We found 10 candidate TFs predicted to recognize differentially the sequence including rs11172113 (log2 Fold Change > 2, **Table S2**). Among these, seven TFs were reported to be expressed in artery tissues from GTEx. We prioritized three TFs that had functions relevant to arterial disease (**Figure 3A**): the methyl CpG binding protein 2 (MECP2) that exhibits a regulatory role in VSMC phenotypic modulation and neointima formation^31, 32^; *SNAI1* that encodes zinc finger protein SNAIL, a member of a family of transcriptional repressors involved in the control of endothelial and epithelial to mesenchymal transitions and fibroblast activation^33, 34^; and RAS-responsive element binding protein 1 (RREB1), a RAS transcriptional effector known to cooperate with TGFβ/SMAD pathway to regulate epithelial to mesenchymal transition^35^. All three TFs were abundantly expressed in SMCs, fibroblasts and endothelial clusters inquired from a single nuclei RNA-Seq dataset of diseased human coronary arteries^36, 37^, with *MECP2* showing higher expression in the SMC cluster, compared to *SNAI1* and *RREB1* (**Figure S5).**

**Figure 3.**
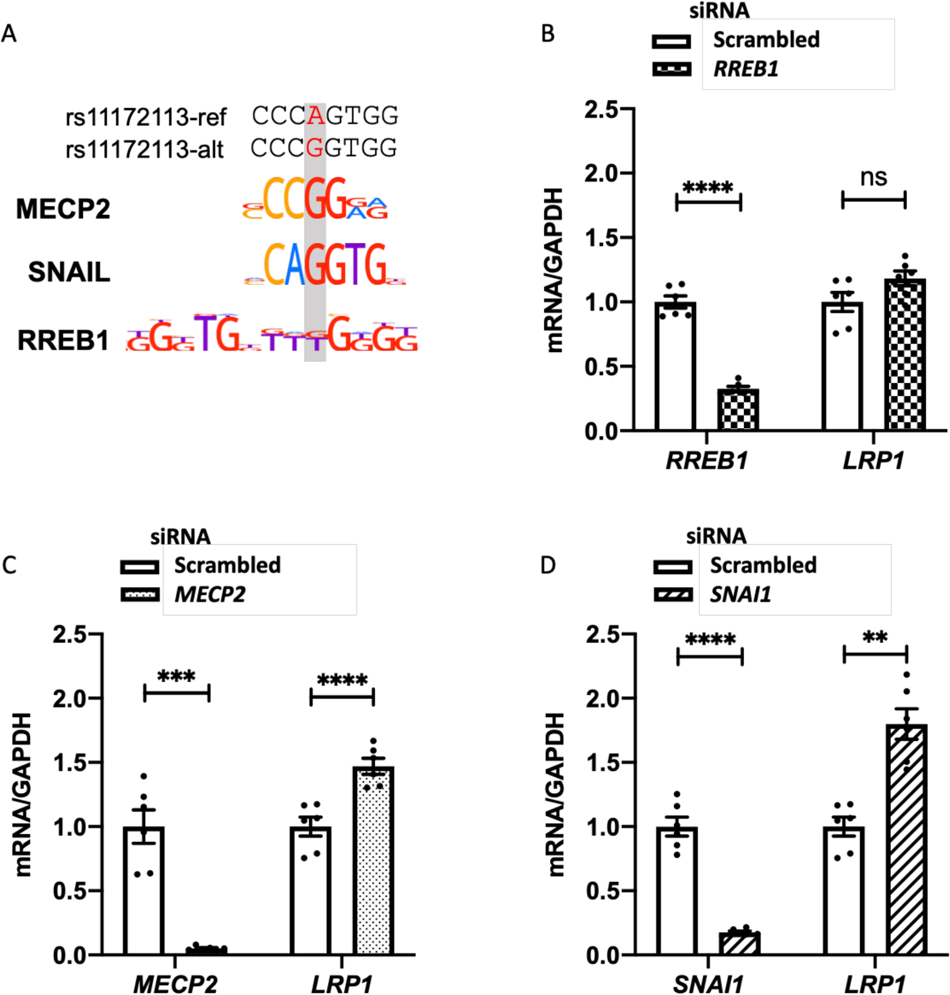
Transcription factors MECP2 and SNAIL are repressors of *LRP1* expression. **A**: Representative images showing alignment of MECP2, SNAIL and RREB1 transcription factors motifs to rs11172113 T or C allele. **B-D**: Bar plots showing mean±SEM of relative RNA levels of *RREB1* (**B**), *MECP2* (**C**), *SNAI1* (**D**) and *LRP1* detected using qPCR following treatment with siRNAs targeting *RREB1* (**B**), *MECP2* (**C**) and *SNAI1* (**D**), compared to scrambled siRNA. Unadjusted T-test: *p<0.05, **p<0.01, ***p<0.001, ****p<10^-4^, ns: not significant.

We found a significant increase in *LRP1* expression when iPSC-derived SMCs (CT genotype for rs11172113) were treated with siRNA targeting *MECP2* and *SNAI1* but not *RREB1* (**Figure 3B-D**), supporting a repression of *LRP1* expression by MECP2 and SNAIL. To determine whether this repression was dependent on rs11172113, we generated homozygous iPSCs for each allele (**Figure 4A, Table S1**). SMCs derived from each genotype presented similar expression levels of *LRP1* (**Figure S6**). However, the knockdown of *MECP2* or *SNAI1* resulted in a significant increase *LRP1* expression only in iPSC-SMCs with the CC genotype (**Figure 4B-C**). Chromatin immunoprecipitation (ChIP) using antibodies for both TFs in iPSC-SMCs with each genotype (CC, CT, or TT) confirmed SNAIL (**Figure 4D)** but not MECP2 (**Figure S7**) to bind to this enhancer only in cells with the CC genotype (**Figure 4B-C**). In summary, these data suggest that rs11172113-C allele is specifically recognized by SNAIL, a repressor of transcription expressed in VSMCs.

**Figure 4.**
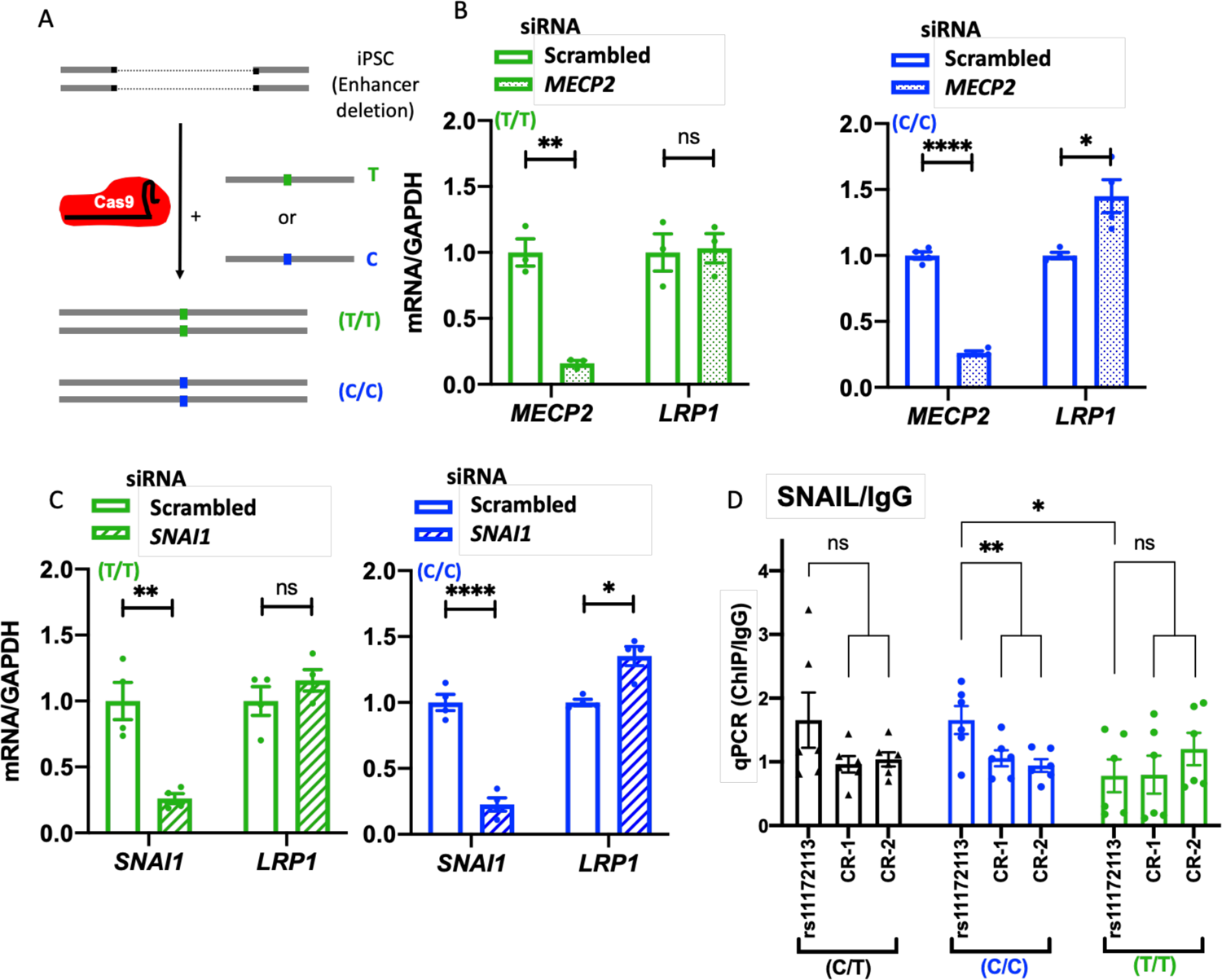
SNAIL selectively binds to rs11172113-C allele. **A**: iPSCs with homozygous TT alleles or CC alleles on rs11172113 were generated by homologous recombination on iPSCs with deletion of rs11172113 enhancer region. **B-C**: Bar plots showing mean±SEM of relative RNA levels of *MECP2* (**B**), *SNAI1* (**C**) and *LRP1* detected using qPCR following treatment with siRNAs targeting *MECP2* (**B**) and *SNAI1* (**C**), compared to scrambled siRNA in iPSC-SMCs harbouring homozygous T (green, left panel) or C allele (blue, right panel) for rs11172113. Unadjusted T-test: *p<0.05, **p<0.01, ***p<0.001, ****p<10^-4^, ns: not significant. **D:** Chromatin immunoprecipitation (ChIP) targeting SNAIL in iPSC-11.10 and homozygous rs11172113 C/C or T/T (two clones each). Immunoprecipitated material was evaluated using qPCR targeting and rs11172113 region and two control regions (no enhancer marks in SMCs or artery tissue). The ratio of SNAIL ChIP to ChIP with rabbit IgG is given. Each clone was assessed in triplicate. Unadjusted T-test: *p<0.05, **p<0.01, ns: not significant.

### *LRP1* KO induces TGFβ activation and extracellular matrix remodeling in iPSC-SMCs

Given the existing conflicting data on LRP1 role in SMCs, we aimed to assess the cellular consequences of LRP1 loss of function in a human iPSC-SMCs model. We introduced a frameshift indel insertion at either exon 2 or exon 5 of *LRP1* using CRISPR/Cas9 (**Figure 5A**) and obtained 4 *LRP1* knockout (KO) iPSC clones with a complete loss of LRP1 protein expression (**Figure S8-9, Table S1**), including after differentiation into SMCs (**Figure 5B**). Compared to WT, we found a decrease in proliferation and migration capacities in *LRP1* KO iPSC-SMCs (**Figures S10-13**). Data in mice described LRP1 as a ligand for intracellular calcium channels in SMCs, promoting their contractile function^21^. However, our *LRP1* KO iPSC-SMCs model had no significant ability to contract a collagen gel lattice or on intracellular calcium release in response to two contractile inducers (carbachol and angiotensin II, **Figures S14-15**).

**Figure 5.**
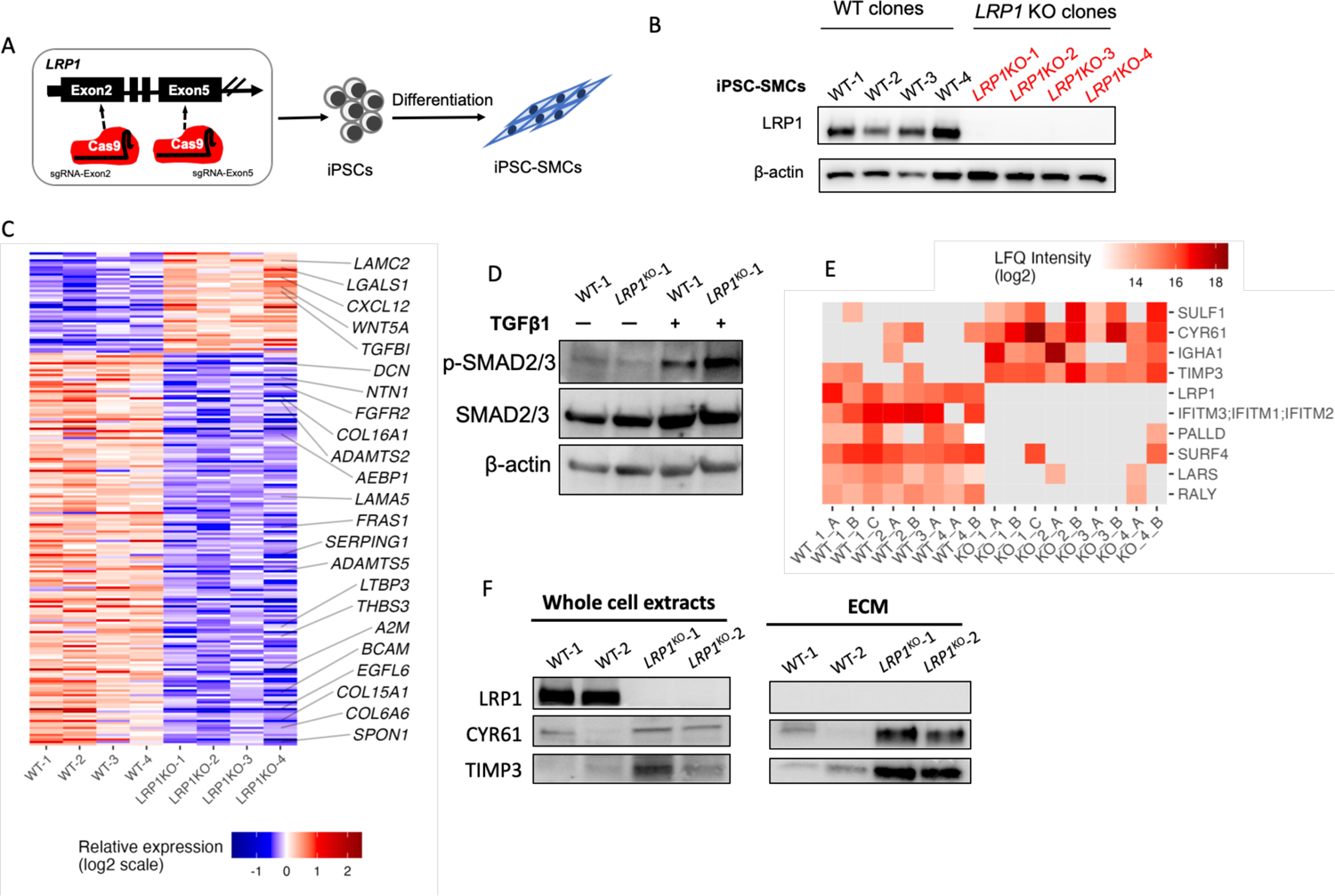
Transcriptomic and proteomic changes following *LRP1* KO in iPSC-SMCs. **A**: Illustration of indel frameshift in exons of *LRP1* to generate knockout iPSCs using CRISPR/Cas9 prior to differentiation into SMCs **B:** Representative Western blot images showing LRP1 protein expression in WT and *LRP1* KO iPSC-SMCs. **C**: Heatmap representation of relative RNA expression for differentially expressed genes between *LRP1* KO and WT iPSC-SMCs. Genes involved in “collagen containing extracellular matrix” (gene ontology 0062023) are indicated **D**: Representative Western blot images showing the expression of phospho-SMAD2/3 (p-SMAD2/3) and total SMAD2/3 (SMAD2/3) in WT and *LRP1* KO iPSC-SMCs in the presence of TGFβ1 protein for 1 hour. **E**: Heatmap representation of label free quantification (LFQ) for proteins over or underrepresented in extracellular extracts of *LRP1* KO and WT iPSC-SMCs. **F**: Representative Western blot images showing the expression of LRP1, CYR61 and TIMP3 in whole cell extracts or decellularized extracts (ECM).

Global gene expression changes using bulk RNA sequencing indicated 213 genes to be differentially expressed between WT and *LRP1* KO cells, with a majority of genes upregulated in the *LRP1* KO cells (**Figure 5C**). Differentially expressed genes were enriched for genes involved in collagen-containing extracellular matrix (*P* value=3.1×10^-11^, **Figure 5C, Figure S16, Table S3**), connective tissue development (*P*-value= 3.2×10^-6^) and lung development (*P*-value= 3.3×10^-6^, **Figure S16, Table S3**). Among genes deregulated in *LRP1* KO, we highlight the transforming growth factor beta induced protein gene (*TGFBI*), an exocrine protein induced by the TGF-β signaling pathway^38^ (**Figure 5C**). Interestingly, we found an increased amount of p-SMAD2/3 in *LRP1* KO SMCs after stimulation by TGF-β1 for 1 hour, and no change in total SMAD2/3 (**Figure 5D**), confirming that canonical TGF-β signaling is enhanced in this model.

The regulation of ECM deposition was the main impaired pathway in our *LRP1* KO iPSC-SMCs, a key pathway in several arterial diseases that is specifically suspected in the case of FMD and SCAD ^39, 40^. To investigate further this mechanism, we assayed changes using label-free quantification (LFQ) with MaxQuant software^41^ of ECM composition by mass spectrometry of decellularized extracts of *LRP1* KO and WT iPSC-SMCs. Despite similar protein content of most abundant ECM proteins (**Figure S17**), several proteins were systematically detected only in one of the two conditions. In particular, *LRP1* KO SMCs secreted increased levels of CYR61, a cysteine-rich angiogenesis inducer and known ligand for LRP1 involved in apoptosis, adhesion, migration and vascular integrity^42^. We also found higher amounts of TIMP3, an inhibitor of the matrix metalloproteinases involved in ECM degradation and known to bind the extracellular domain of LRP1 to facilitate the clearance of target metalloproteinases (**Figure 5E**). *CYR61* and *TIMP3* were confirmed to be abundantly expressed in SMC, fibroblasts and endothelial clusters inquired from single nuclei RNA-Seq dataset of diseased human coronary arteries^36^ (**Figure S6**). Immunoblotting on whole cell or ECM extracts generated from iPSC-SMCs confirmed the presence of increased amounts of CYR61 and TIMP3 in *LRP1* KO SMCs, compared to WT cells (**Figure 5E**), which we confirmed in iPSC-SMCs after LRP1 knockdown for 3 days (**Figure S18**).

## Discussion

In this integrated study, we provide molecular and cellular mechanisms underlying the genetic association in *LRP1* locus involved in the risk for multiple major vascular diseases. Through conditional analyses and colocalization of association signals, we pointed at the intronic variant rs11172113 as the most likely causal variant for all associations reported. *In vitro* experiments conducted on several genome-edited iPSC-derived SMCs models support rs11172113 to be located in an enhancer region that controls the expression of *LRP1* through the action of transcription factors MECP2 and SNAIL in an allele-specific repression mechanism. Our results also support *LRP1* depletion to decrease iPSC-derived SMCs proliferation and migration, potentiate canonical TGF-β signaling and lead to partial ECM remodeling involving enhanced secretion of CYR61 and TIMP3 proteins.

Our results illustrate the complexity of regulatory mechanisms at stake at GWAS loci involving potentially multiple target genes. Following the deletion of a 100bp region centered on the most probable causal variant in this locus in iPSC-derived SMCs, we found a significant decrease in the expression of 2 genes: *LRP1* and *NAB2*. This finding suggests that both genes may share this enhancer region, at least in our iPSC-SMCs models. NAB2 protein has been reported as a transcriptional corepressor able to bind early growth response protein 1 to regulate proliferation, migration and survival of vascular cells in response to vascular injury^43, 44^, including in the context of atherosclerosis and hypertension^45^. Although we cannot exclude the possibility of more than one target gene in this locus, which may explain its high genetic pleiotropy, genetic colocalization of eQTL signals in arterial tissues with GWAS signals only supported *LRP1* as the most likely causal gene at the locus. Future studies involving potentially additional cell types, relevant to vascular injury for instance, are needed to address this discrepancy.

Using *in silico* prediction and gene knockdown in genome-edited cells, we found that transcription factors SNAIL and MECP2 repress *LRP1* expression specifically in the presence of the C allele of rs11172113. Through ChIP experiments, we confirmed *in silico* predictions and showed that SNAIL directly binds to the rs11172113 region in an allelic specific manner. SNAIL is a master regulator of endothelial to mesenchymal transition involved in tumor metastasis, kidney fibrosis and pulmonary hypertension^46–48^. Although the function of SNAIL in vascular smooth muscle cells is not known, it was recently found that closely related transcription factor SLUG was involved in SMC phenotypic regulation during atherosclerosis^49^. The absence of interaction with MECP2 could be due to technical limitations of the ChIP approach, in particular, the limited availability of ChIP grade antibodies. This, our result does not rule out a potential role of this TF in LRP1 regulation, as it was previously shown to regulate vascular SMC phenotypic plasticity and neointima formation^31, 32^.

Our findings suggest that the functional consequence of rs11172113 allelic variation could be at play when vascular SMCs and fibroblasts undergo major phenotypic changes, such as wound repair, neointima formation, arterial remodeling and atherosclerosis, as reported from several *in vivo* studies^19, 21, 22, 50^. Conversely, a lack of *LRP1* repression following arterial injury would be expected to impair arterial contractility and wound repair, two suspected mechanisms in FMD and migraine pathogenesis^11, 51^. Of note, we did not observe rs11172113 allele-specific differences in *LRP1* expression between iPSC-derived SMCs in the absence of TF knockdown. Potential compensatory mechanisms may buffer the expression of *LRP1*, especially in the absence of the phenotypic transitions that cannot be modelled in our *in vitro* modelling, which may potentially involve additional TFs with affinity to this enhancer region.

LRP1 is a multifunctional protein that recognizes hundreds of ligands and functions in a variety of biological processes, including lipid metabolism, endocytosis and signaling pathway regulation^18, 52^. Due to this versatility, the phenotypic effect of LRP1 deletion is highly model-dependent and limited extrapolation is warranted from *in vitro* studies. However, through an exploration of molecular changes in *LRP1* KO iPSC-SMCs, we observed several alterations particularly relevant to the pathogenesis of FMD, arterial dissection, migraine and lung function. We found that LRP1 deficiency led an alteration of the extracellular matrix both in terms of gene transcription and protein content, a finding consistent with a recent study in mice^53^. The hyperactivation of canonical TGFβ signaling in LRP1 deficient cells may explain changes in gene expression, given the role of this pathway in promoting fibrosis provided from *in vivo* models^50, 54^. In support of this mechanism, we mention the enrichment for rare mutations in genes from the TGFβ pathway among SCAD patients^55^, the recent report of *FBN1*, known reservoir for TGFβ, as a potential target gene in a SCAD risk locus^12, 56^, and of *TGFBR2* in migraine^57^. Thus, increased expression of *LRP1* in arteries, as genetically predicted in SCAD, FMD and migraine in *LRP1* allele risk carriers, could prelude the dysregulation of TGFβ signaling.

Our study also identified an increased deposition of TIMP3 and CYR61 in the extracellular matrix generated by LRP1 KO SMCs. TIMP3 promotes the degradation and clearance of matrix metalloproteinases, and is suspected to be involved in a wide variety of cardiovascular diseases^58^. LRP1 regulation of TIMP3 endocytosis is well documented in a variety of models^58^. In addition, *TIMP3* was recently prioritized in a SCAD GWAS, which supports a potential mechanistic link between LRP1 and TIMP3 in the context of this arterial disease^56^. On the other hand, CYR61 is a paralogue of connective tissue growth factor (CTGF), reported to be regulated by *LRP1* in arteries.^20, 50^ CYR61 plays an important role in wound healing and was associated to several lung diseases, including chronic obstructive pulmonary disease and pulmonary hypertension,^59, 60^ consistent with *LRP1* association with FEV/FEC. However, the relevance of this mechanism in the other vascular diseases needs further investigation.

Our work presents several limitations. First, iPSC derived SMCs are a useful model but do not fully recapitulate all complex phenotypes of SMCs *in vivo*, which warrants further confirmation of the proposed regulatory mechanisms. Second, we did not prioritize the investigation of the function of *NAB2* in SMCs based on non-consistency of the existing eQTLs data from human arteries, compared to *LRP1*. Thus, we cannot exclude the role of this gene in the physiopathology of vascular diseases with genetic association reportd in this locus. Third, although we showed that TFs MECP2 and SNAIL regulate the expression of *LRP1* specifically through rs11172113 C-allele, we did not demonstrate direct binding of MECP2 to rs11172113 associated enhancer, and additional TFs predicted *in silico* to change their ability to bind to this genomic enhancer may also play a regulatory role. Fourth, our study design does not cover the highly suspected multiple cell types interactions involving endothelial cells and fibroblasts to fully depict the functional translation the genetic association with such a large panel of arterial diseases.

In conclusion, we provide a comprehensive functional investigation of a highly pleiotropic genetic locus involved in the genetic susceptibility to several vascular diseases, and propose a plausible molecular and cellular mechanisms linking the most likely causal variant to gene expression regulation of *LRP1*. We provide evidence for an important impact of the most likely target gene, *LRP1*, on the regulation of proliferation and migration. Based on the exploration of ECM synthesis capacity of several *in vitro* iPSC-SMCs models, we propose a mechanism involving, at least partially, potentiated activity of the canonical TGFβ signaling. Our work provides genetically-driven biological mechanisms to elucidate the pathogenesis of several important and under-studied cardiovascular diseases.

## Supporting information

Supplemental methods and results

## Acknowledgements

This work benefited from expertise and support from the iPSC technical facility at PARCC, and the high throughput sequencing core facility of I2BC (Centre de Recherche de Gif – http://www.i2bc.paris-saclay.fr/). The Genotype-Tissue Expression (GTEx) Project was supported by the Common Fund of the Office of the Director of the National Institutes of Health, and by National Cancer Institute (NCI), National Human Genome Research Institute (NHGRI), National Heart, Lung, and Blood Institute (NHLBI), National Institute on Drug Abuse (NIDA), National Institute of Mental Health (NIMH), and National Institute of Neurological Disorders and Stroke (NINDS).

## Methods

### Conditional analysis

We applied the GCTA COJO method^26^ to perform local stepwise model procedure to select independently associated SNPs at the *LRP1* locus. We used published GWAS summary statistics for migraine and FMD^11, 61^. The local LD structure was provided in a reference file from the haplotype reference consortium (http://www.haplotype-reference-consortium.org/site). LocusZoom (http://locuszoom.org/) was used for visualization.

### Colocalization analyses with diseases and traits

Summary statistics were retrieved from individual studies^11, 14, 61, 62^. The approximate Bayes factor colocalization analyses were performed for 4 traits or diseases for which genome-wide significant association in *LRP1* was reported (Migraine, FMD, pulse pressure and FEV1/FVC). Signal colocalization was evaluated using R coloc package (v5.1.0) with default values as priors^63^. The H4 coefficient indicating the probability of the two traits to share a causal variant was reported. We queried the GTEx database (v8 release) 61 with lead variants in 3 arterial tissues for *LRP1* (permutations q value < 0.05). For colocalization, all SNP-gene associations in the 3 arterial tissues were retrieved from GTEx to compare with the diseases summary statistics. Multitrait colocalization to prioritize causal variant was performed using HyPrColoc package.^27^

### Cell culture

Rat SMCs (A7r5) were purchased from ATCC® (Manassas, USA) and maintained in DMEM supplemented with 10% FBS (Thermo Fisher Scientific, Waltham, MA, USA). Human iPSC line SKiPS-31.3 was obtained by reprogramming of human dermal fibroblast of a healthy male adult volunteer as previously described^64^. iPS cell lines 11.10 and 12.10 were purchased from Cell Applications (San Diego, CA) and were identified as female using *SRY* PCR. All iPSC lines were maintained in mTeSR^TM^ Plus medium (STEMCell Technologies). iPSCs were differentiated in 24 days into mesoderm-derived vascular SMCs using a previously described protocol, detailed in the supplemental material^29^. After day 24, iPSC-SMCs were maintained in DMEM supplemented with 5 µg/ml insulin, 0.5 ng/ml EGF, 2 ng/ml bFGF, 5% FBS and 1% penicillin-streptomycin (all from Thermo Fisher Scientific, Waltham, MA, USA).

### Dual-luciferase enhancer reporter assay

To generate pGL-TK minimal promoter reporter plasmid, TK-minimal promoter was excised from pRL-TK (Promega, Madison, Wisconsin, USA) using BglII/HindIII restriction enzymes and instered into corresponding sites of pGL4.12 (Promega, Madison, Wisconsin, USA). A 902bp DNA fragment centered on rs11172113) was amplified from genomic DNA of iPSC-11.10 (primers: F: tggccggtaccTCAGAAGGAAGGAGGGAGGT, R: ctcgaggctagcCATTCTCACCCCACTTCCCC), and inserted between KpnI and NheI sites of pGL-TK. For reporter assay, cells were seeded at a density of 5,000 cells per well in 96-immuno white plates (Thermo Fisher Scientific, Waltham, MA, USA). The pGL-TK derived plasmids (firefly luciferase) and pRL-TK (Renilla luciferase) vector were co-transfected into cells in a 4:1 ratio. The A7r5 cells and iPSC-SMCs (from 31.3 iPSC clone) were transfected respectively using FuGENE HD Transfection Reagent Complex (Promega, Madison, Wisconsin, USA) and Amaxa Basic Nucleofector kit for primary mammalian SMCs (LONZA, Basel, Switzerland) using Nucleofector 2b device (LONZA, Basel, Switzerland) according to manufacturer’s instructions. 48h after transfection, the Firefly and Renilla luciferase were measured by performing Dual-Luciferase Reporter Assay System (Promega, Madison, Wisconsin, USA), following manufacturer’s instructions. Signal was recorded using Mithras LB 940 Multimode Microplate Reader machine (Berthold Technologies, Bad Wildbad, DE). The enhancer capacity was calculated by the ratio of Firefly to Renilla luciferase. Statistical significance was evaluated using Student’s two sample t-test with unequal variances.

### Genome editing in iPSCs by CRISPR/Cas9

Guide RNAs (gRNA) were designed using the CRISPOR online tool (http://crispor.tefor.net/)^65^. The following sequences were targeted, with the indicated predicted cut sites: rs11172113-141-fw (CAAAGCAGAGGCCCAGACTC*AGG*) targeting position 45bp upstream of rs11172113, rs11172113-218-rev (CCTCCCAAACCCAAACCGAG*AGG*) targeting position 42bp downstream of rs11172113, LRP1-159-rev (GGGCCTCGTCAGATCCGTCT*GGG*), targeting position 165 of *LRP1* cDNA, LRP1-547-rev (GCCTTGCAGGAGCGGTTATC*CGG*), targeting position 553 of *LRP1* cDNA and rs11172113-141/218-del (GAGGCCCA<u>GAGG</u>TTTGGGTT*TGG*) targeting the 88bp deletion site generated around rs11172113 (junction is underlined). Designed oligos were inserted into plasmids pSpCas9(BB)-2A-Puro (PX459) V2.0 or pSpCas9(BB)-2A-Hygro, which were gifts from Feng Zhang and Ken-Ichi Takemaru (Addgene plasmids # 62988 and # 127763).^66^

Induced pluripotent stem cells were transfected with a total of 5 μg target Cas9/gRNA plasmids with or without 1 μg repair templates using Human Stem Cell Nucleofector Kit (LONZA, Basel, Switzerland) and Nucleofector 2b device (LONZA, Basel, Switzerland), following manufacturer’s instructions. Medium was changed 48h after transfection to mTeSR^TM^ Plus medium containing with 0.5 μg/ml puromycin and/or 25 μg/ml hygromycin for 2 days to select cells transfected with pSpCas9, after what they were grown with unsupplemented mTeSR^TM^ Plus medium. To delete 88 base pairs of enhancer sequence centered on rs11172113, pSpCas9-rs11172113-141-fw-puro and pSpCas9-rs11172113-218-rev-hygro were co-transfected into iPSCs (11.10 cell line). To obtain *LRP1* knock-out cells, pSpCas9-LRP1-159-rev-puro or pSpCas9 LRP1-547-rev-puro were transfected in iPSC (11.10 cell line, separate experiments). To obtain clones homozygous for rs11172113, iPSC with enhancer deletion (clone rs11172113-141/218-10) were co-transfected with pSpCas9-rs11172113-141/218-del-puro and a 872bp PCR product amplified from pGL-TK-rs11172113-C or pGL-TK-rs11172113-T (primers: F: CTGGCAGAACTTCCTCCTGA, R: GAAGTGGGGTGAGAATGCCT). The genomic DNA of selected iPSCs clones was extracted using QIAamp DNA Micro Kit (Qiagen, Hilden, Germany). All sequences of enhancer deletion or allele insertion were verified using Sanger sequencing.

### Knockdown

Cells were seeded in 6-well plates at a density of 200,000 cells per well to reach ∼80% confluence. Cells were transfected with Silencer® Select N°1 negative control siRNA or Silencer® Select predesigned siRNA for *MECP2* (s8644), *SNAI1* (s13185) or *RREB1* (s12355) (Thermo Fisher Scientific, Waltham, MA, USA) using Opti-MEM and Lipofectamine RNAiMAX transfection reagent (Invitrogen, Waltham, Massachusetts, USA) for 48 hours.

### Antibodies

Following antibodies were used in Western blot experiments: rabbit anti-LRP1 (ab92544, Abcam, Cambridge, UK), mouse anti-β-actin (sc-47778, Santa Cruz Biotechnology, Dallas, Texas, USA), rabbit anti-phospho-SMAD2/3 (#8828) and anti-SMAD2/3 (#3102, Cell Signaling Technology, Danvers, Massachusetts, USA), mouse anti-TIMP3 (LS-B2576, Lifespan Biosciences, Seattle, Washington, USA), mouse anti-CYR61 (sc-374129, Santa Cruz Biotechnology, Dallas, Texas, USA). Following antibodies were used for ChIP: mouse anti-SNAI1 (G7) (sc-271977, Santa Cruz Biotechnology, Dallas, Texas, USA), rabbit anti-MECP2 (ab195393, Abcam, Cambridge, UK).

### Chromatin immunoprecipitation

Chromatin immunoprecipitation was performed as previously described ^67^. Briefly, 10^7^ cells (1 confluent 150mm dish) were crosslinked with 1.1% Formaldehyde for 10 minutes. Reaction was quenched with 2.5M Glycine and cells were washed, collected in cold PBS, pelleted and flash frozen. After two mild lysis steps, cells were lysed in lysis buffer C (1 mM EDTA, 0.5 mM EGTA, 10 mM pH7.5 Tris-HCl, 100 mM NaCl, 0.1% Na-Deoxycholate, 0.5% N-lauryl Sarcosine and protease inhibitor cocktail), and chromatin was sonicated using on EpiShear Porbe Sonicator (Active Motif, Carlsbad, CA, USA). Sheared chromatin was incubated overnight with 100µL Protein G Dynabeads (Thermo Fisher Scientific, Waltham, MA, USA) complexed with 10µg antibody or rabbit IgG. Precipitated DNA was de-crosslinked overnight at 65°C and purified using Phenol/Chloroform extraction. Details for the method are given in the Supplemental material.

### Cellular assays

Cellular assays including cell viability assay, EdU flow cytometry assay, wound healing assay, collagen matrix contraction assay and intracellular calcium measurement were performed as previously described.^68^ Details for each method are given in the Supplemental material.

### Statistical analyses

Statistical assessment of RNA-Seq were performed using DESeq2 (v1.32.0) package in R (v4.1.0).^69, 70^ Differential detection analysis in mass spectrometry experiments was done using DEqMS package (v1.10.0) and is detailed in the Supplemental material.^41, 71–73^ For all other experiments, statistical significance was evaluated using Student’s two sample t-test with unequal variances. Further details are given in the Supplemental material.

