## Supplemental methods and results for "Regulatory mechanisms in multiple vascular diseases locus *LRP1* involve repression by SNAIL and extracellular matrix remodeling"

### Supplemental material

**Detailed methods** p2

**Supplementary Tables S1-S3** p9

**Supplementary Figures S1-S18** p12

**References** p32

### Detailed Methods

#### iPSC differentiation into SMCs

iPSCs were differentiated in 24 days into SMCs using a small molecule-based monolayer differentiation protocol[1]. In brief, iPSC monolayers of 85% cell confluence were dissociated into single cells by Accutase (Thermo Fisher Scientific), seeded on Matrigel-coated dish at $10^5$ cells/$cm^2$ and cultured for 36 hours in E8BAC medium (E5 medium[2] plus 5 ng/ml BMP4, 25 ng/ml Activin A, 19.4 mg/l insulin, 10 μM Y27632 and 1 μM CHIR99021). Cells were dissociated and seeded on a new dish coating Matrigel at 1.6 x $10^4$ cells/$cm^2$ in E6T medium (E5 medium supplemented with 19.4 mg/l insulin, 1.7 ng/ml TGF-β1 and 10 μM Y27632) for 18 hours and then cultured with E5F medium (E5 medium supplemented with 19.4 mg/l insulin and 100 ng/ml FGF2) (day 3-7). From day 8 to day 11, cells were treated with FVR medium (E5 medium supplemented with 19.4 mg/l insulin, 50 ng/ml VEGF and 5 μM RESV) and changed every other day. From day 12, cells were cultured in E6-TP medium (E5 medium supplemented with 19.4 mg/l insulin, 5 μM RESV, 10 ng/ml PDGFbb and 1.7 ng/ml TGF-β1). On day 16, cells were cryopreserved when necessary or split in a new Matrigel-coated dish at $1x10^5$ cells/$cm^2$ cell density. After day 24, iPSC-SMCs were maintained in DMEM supplemented with 5 μg/ml insulin (Thermo Fisher Scientific, Waltham, MA, USA9), 0.5 ng/ml EGF (Thermo Fisher Scientific, Waltham, MA, USA), 2 ng/ml FGF (Thermo Fisher Scientific, Waltham, MA, USA) and 5% FBS. 5% FBS and 1% penicillin-streptomycin (Thermo Fisher Scientific, Waltham, MA, USA). The cells were passaged with 0.25% Trypsin/EDTA solution (Thermo Fisher Scientific, Waltham, MA, USA).

#### Quantitative RT-PCR analysis

Cells were collected and washed with PBS once. The purification of total RNA was performed using RNeasy Plus Mini kit (Qiagen, Hilden, Germany) following instructions of the manufacturer. The complementary DNA synthesis was performed by reverse transcription with 1 μg of total RNA using the iScript cDNA Synthesis Kit (BIO-RAD, Hercules, California, USA). Quantitative real-time PCR was performed using SYBR reagent (GoTaq qPCR system, Promega, Madison, Wisconsin, USA) and primers designed with online tool (https://primer3.ut.ee/), and the reaction was run on StepOnePlus machine from Applied biosystems. Relative gene expression quantities were determined by comparative $C_T$ methods in StepOne software and were normalized to housekeeping genes *GAPDH*, *ACTB* and *SDHA*.

Each reaction was run at least in triplicates and statistical significance was evaluated using Student's two sample t-test with unequal variances.

**Chromatin Immunoprecipitation-qPCR**

The chromatin immunoprecipitation was performed following a previously described protocol [3]. Cells (one 150mm dish at 100% confluency, ~$10^7$ cells) were incubated with 10% cross-linking solution (11% formaldehyde, 50 mM HEPES pH=8, 100 mM NaCl, 1mM EDTA and 0.5mM EGTA) for 10 mins at room temperature, and we then added 5% 2.5 M glycine to quench formaldehyde. The cells were washed twice by PBS, harvested and flash-frozen in liquid nitrogen.

The cells were lysed in three steps using first lysis buffer A (50 mM pH 7.5 HEPES-KOH, 140 mM NaCl, 1 mM EDTA, 10% glycerol, 0.5% NP-40, 0.25% Triton X-100 and protease inhibitor cocktail) rocking 10 mins at 4°C. Cells were the resuspended in lysis buffer B (200 mM NaCl, 1 mM EDTA, 0.5 mM EGTA, 10 mM pH 7.5 Tris and protease inhibitor cocktail) and rocked 10 mins at 4°C. Finally cells were resuspended in 1 mL lysis buffer C (1 mM EDTA, 0.5 mM EGTA, 10 mM pH7.5 Tris-HCl, 100 mM NaCl, 0.1% Na-Deoxycholate, 0.5% N-lauryl Sarcosine and protease inhibitor cocktail). The suspension was sonicated using a EpiShear Probe Sonicator (Active Motif, Carlsbad, CA, USA), for a total of 20 mins in cycles of 30 seconds spaced by a 40 seconds rest time with 120 W, 20 kHz, 80% Amplitude.

The Protein G Dynabeads magnetic beads (100 µl/reaction) were washed with 0.5% BSA/PBS twice and incubated and rotated at 4°C overnight with 10 µg rabbit IgG (Invitrogen, Waltham, Massachusetts, USA) or antibodies: anti-SNAI1 (G7) (Santa Cruz Biotechnology, Dallas, Texas, USA), anti-MECP2 (Abcam, Cambridge, UK). Then the beads-antibody complexes were washed 3 times and resuspended in 100 ul 0.5% BSA/PBS.

The sonication products were incubated with beads-antibody products rocking overnight at 4°C. Then the IP reactions were washed 4-6 times by wash buffer (50 mM pH 7.6 HEPES, 1 mM EDTA, 0.7% Na deoxycholate, 1%NP-40 and 0.5 M LiCl) and once with TBS. Then the products were eluted by elution buffer (50 mM pH8 Tris, 10mM EDTA and 1% SDS) at 65°C for 10-15 min and then were spined 15000g for 1 min. The supernatant was incubated at 65°C overnight to reverse cross-linking. The samples were incubated with RNase A (0.2 µg/µL) for 1-2 hours at 37°C and then add proteinase K (final concentration is 0.2) to incubated at 55°C for 1-2 hours. The DNA was purified by Phenol/Chloroform method and was resuspended in 60 µL 10mM pH 8 Tris-HCl. The DNA concentration was measured using Qubit HS DNA assay. The quantitative RT-PCR analysis was performed as mentioned above. Following primer pairs were used: rs11172113 (F: CACCCTGTCTGTCTGTCTGT, R: CCCAGTGGCTCTTTCCTGA), Control 1 (F: AGCGAGGTTTATGGTGGACA, R: AGATGTTGGGCTGAAGTGGA), Control 2 (F: AACCTGAGGAGAGTGCAGAC, R: TTCCCCACCTGACTTCTCAC).

**Western blotting**

After cells were washed once by PBS, 100 μl 2 x Laemmli Sample Buffer (BIO-RAD, Hercules, California, USA) was added for 6-well plates. Samples were collected and incubated at 95 °C for 10 minutes supplemented with 5% β-Mercaptoethanol. Then samples were centrifuged for 5 minutes at max speed and the supernatant was transferred to a fresh 1.5mL tube. The protein samples were processed to SDS-PAGE and transferred to nitrocellulose membranes. After blocking for 1 hour at room temperature in TBST buffer supplemented 5% (w/v) BSA, membranes were incubated at 4°C overnight with agitation following primary antibodies (1:1000 in 2% BSA buffer): anti-LRP1 (Abcam, Cambridge, UK), anti-β-actin (Santa Cruz Biotechnology, Dallas, Texas, USA), anti-phospho-SMAD2/3 (Cell Signaling Technology, Danvers, Massachusetts, USA), anti-SMAD2/3 (Cell Signaling Technology, Danvers, Massachusetts, USA). Membranes were then rinsed three times for 10 minutes each with TBST and incubated with HRP conjugated secondary antibodies (BIO-RAD, Hercules, California, USA) diluted 1:20000. After three times rinsing for 10 minutes with TBST, membranes were incubated with SuperSignal West Pico PLUS Chemiluminescent substrate (Thermo Fisher Scientific, Waltham, MA, USA) and visualized by FujiFilm LAS-4000 mini system (FUJIFIML, Minato City, Tokyo, Japan).

**Cell viability assay**

Cells (5,000 cells/well) were seeded in 96-immuno white immune plates (Thermo Fisher Scientific, Waltham, MA, USA) with 8 duplicates. CellTiter-Glo 2.0 assay (Promega, Madison, Wisconsin, USA) was used to measure cell viability. Plates were loaded with CellTiter-Glo reagents and the luminescent signal was recorded using Mithras LB 940 Multimode Microplate Reader machine. Measurement of luciferase was performed daily for 5 days, beginning from the 4$^{th}$ hour after cell seeding (4 h). The viability of all luciferase measurements was normalized to the values at 4 h. Statistical significance was evaluated using Student's two sample t-test with unequal variances.

**EdU flow cytometry assay**

Cells were seeded at a density of 150,000 cells (6-well plate) one day before. Cells were incubated with 10 μM EdU (5-ethynyl-2'-deoxyuridine, Life technologies) for 4 hours and then harvested to perform Click-iT reaction by using Click-iT EdU flow cytometry Alexa Fluor 488 assay kit (Life technologies, Carlsbad, CA, USA). Fluorescence measurements were performed by using LSRfortessa X-20 (BD Biosciences, New Jersey, USA). FACS results were analyzed by Flowjo (BD Biosciences, New Jersey, USA). Statistical significance was processed using Student's two sample t-test with unequal variances.

**Transwell assay**

100 μl of cell suspensions (40,000 cells/ml) were plated on the top of filter membrane in the Transwell insert (24-well Transwell chamber with 8 μm pore size, Corning, New York, USA) and incubated for 10 mins at 37 °C to precipitate cells to bottom. 600 μl medium were added into the bottom chamber. Chambers were incubated under the cell culture condition for 20 hours. The inserts then were collected, fixed in 70% ethanol and dyed with 0.2% crystal violet for 15 min. The migrated cells were imaged and counted using ImageJ tool. Statistical significance was assessed using Student's two sample t-test with unequal variances.

**Wound healing assay**

Cells were seeded one day before to reach confluent monolayers (24-well plates). Straight wounds were scratched through the entire center of the well by a 10-μl pipette tip. The plates were then washed with media once and photographed for 12 hours with 1-hour intervals. Three different fields at each time point on each plate were imaged and the wound area was measured by Image J. The wound closure was calculated by the wound area in each period as a percentage of the initial wound area at 0 h. Statistical significance was evaluated using Student's two sample t-test with unequal variances.

**Collagen contraction assay**

Cells at a density of $2.5\times10^5$/well were harvested and resuspended in culture medium. Collagen gel was prepared by mixing cell suspension and collagen solution and seeded into 48-well plate with Cytoselect Cell contraction Assay Kit (CELL BIOLABS Inc, San Diego, CA, USA) according to manufacturer's instructions. Gels were imaged 20 h post-seeding using a RICOH MP3503 scanner (RICOH, Tokyo, Japan). Collagen gel areas were determined using Image J software. Gel area changes were calculated by the percentage of gel area for each condition compared to the gel area of the control (without cells). Statistical significance was evaluated using Student's two sample t-test with unequal variances.

**Intracellular calcium influx measurements**

Cells were seeded overnight in 96-well black/clear bottom plates coated with Matrigel at the density of 40,000 cells per well. The washing buffer was supplied with 5 mM HEPES and 250 mM probenecide in HBSS with calcium and prepared in the same day. Cells were washed twice and incubated with calcium-sensitive dye Fluo-4-AM (2 μM) (Ex/Em=480 nm/525 nm) for 1 h at 37 ºC avoiding exposition to light. Cells were washed twice and incubated in washing solution for 20 mins at room temperature in the dark. The infusion of vasoconstrictor solution carbachol was performed by PC-controlled pump. The fluorescence was detected every 1.52 second in a 180 second range. Fluorescence measurements were normalized to the average of first 10 signals. Statistical significance was evaluated using Student's two sample t-test with unequal variances.

**RNA-Seq and analysis**

Isolated total RNAs were quantified using Qubit ds DNA HS assay kit (Thermo Fisher Scientific, Waltham, MA, USA). PolyA+ mRNA libraries were prepared from 200ng-1μg of total RNA using QIAseq Stranded RNA library preparation kit (Qiagen) according to manufacturer's instruction. Libraries were sequenced on a NextSeq500 instrument (Illumina, San diego, CA, USA) using NextSeq 500/550 High Output Kit v2 (75 cycles). Reads were demultiplexed using bcl2fastq2 (v2.18.12) and adapters were trimmed using Cutadapt (v1.15). Reads were then mapped to human genome (GRCh38.104) using STAR aligner (v2.7.9a) with following options: --outFilterType BySJout --outFilterMultimapNmax 20 --alignSJoverhangMin 8 --alignSJDBoverhangMin 1 --outFilterMismatchNmax 999 --outFilterMismatchNoverReadLmax 0.04 --alignIntronMin 20 --alignIntronMax 1000000 --alignMatesGapMax 1000000. We used per gene read counts as direct input for differential expression analysis using DESeq2 (v1.32.0) package in R (v4.1.0), keeping only genes with mean read count over 1[4]. We transformed the count matrix using variance stabilizing transformation with option blind=FALSE. Differentially expressed genes were determined using res function and log2 fold changes were determined using lfcShrink function with ashr shrinkage estimator[5]. Gene ontology term enrichments were calculated using clusterProfiler package (v4.0.5).

**Extracellular matrix collection and protein quantification**

Cells were seeded in 10-cm uncoated dishes for 48-72 h until 100% confluence was achieved. Cells were then rinsed and incubated in the absence of serum for 1h at 37°C. Cells were washed once by PBS and incubated with 20 mM Ammonium hydroxide at room temperature for 5 minutes with gentle agitation to remove all cells. Dishes were washed by copious volume of de-ionized $H_2O$ four times. The matrix was scraped by 0.1% SDS and frozen in -80°C.

For mass spectrometry analysis, sample were first dried, and proteins were denatured in SDS 2% and TEAB 200 mM pH8.5 while disulfide bridges were reduced using TCEP (tris(2-carboxyethyl)phosphine) 10 mM and subsequent free thiols groups were protected using chloroacetamide 50 mM for 5 min at 95°C. Proteins were trypsin-digested overnight using the suspension trapping S-Trap method (Protifi, Farmingdale, NY, USA) to collect peptides as previously described[6]. Eluted peptides were vacuum-dried in a Speed Vac (Eppendorf). LC-MS analyses were performed on a Dionex U3000 RSLC nano-LC system (Thermo Fisher Scientific, Waltham, MA, USA) coupled to a tims-TOF Pro mass spectrometer (Bruker Daltonics GmbH, Bremen, Germany). After drying, digested samples were solubilized in 10 µL 0.1% TFA containing 10% acetonitrile (ACN). One microliter was loaded, concentrated, and washed for 3 min on a C18 reverse-phase precolumn (3 µm particle size, 100 Å pore size, 75 µm inner diameter, 2 cm in length, from Thermo Fisher Scientific, Waltham, MA, USA). Peptides were separated on an Aurora C18 reverse-phase resin (1.6 µm particle size, 100 Å pore size, 75 µm inner diameter, 25 cm in length), connected to a Captive nanoSpray Ionization module (IonOpticks, Middle Camberwell, Australia) with a 60 min run time and a gradient ranging from 99% solvent A, containing 0.1% formic acid in milliQ-grade H2O, to 40% solvent B, containing 80% acetonitrile and 0.085% formic acid in mQH2O. The mass spectrometer acquired data throughout the elution process and operated in the DDA PASEF mode, with a 1.9 s/cycle, with the timed ion mobility spectrometry (TIMS) mode enabled and a data-dependent scheme with full MS scans in PASEF mode. This enabled recurrent loop analysis of a maximum of the 120 most intense nLC-eluting peptides, which were CID fragmented between each full scan every 1.9 s. The ion accumulation and ramp times in the dual TIMS analyzer were set to 166 ms each, and the ion mobility range was set from 1/K0 = 0.6 vs. cm−2 to 1.6 vs. cm−2. Precursor ions for MS/MS analysis were isolated in positive mode with the PASEF mode set to « on » in the 100–1700 m/z range by synchronizing quadrupole switching events with the precursor elution profile from the TIMS device. The cycle duty time was set to 100%, accommodating as many MSMS in the PASEF frame as possible. Singly charged precursor ions were excluded from the TIMS stage by tuning the TIMS using TIMS control software (Bruker Daltonics GmbH, Bremen, Germany).

Identifications (protein hits) and quantifications were performed by comparison of experimental peak lists with a database of theoretical sequences using MaxQuant version 1.6.2.3[7]. The databases used were the Human sequences from the NCBInr database (release March 2020) and a list of in-house frequent contaminant sequences. The cleavage specificity was trypsin's with maximum 2 missed cleavages. Carbamidomethylation of cysteines was set as constant modification, whereas acetylation of the protein N terminus and oxidation of methionines were set as variable modifications. The false discovery rate was kept below 5% on both peptides and proteins.

For differential protein detection, proteins quantified in at least two samples of each condition were kept for further analysis. Label-free quantification intensities from MaxQuant were log2 transformed and normalized using quantile normalization[8]. We performed differential detection analysis using DEqMS package (v1.10.0)[9]. Median peptide counts were used to categorize proteins to compute the variance model. To account for proteins detected only in one situation, protein groups detected in at least 7 WT or KO samples and at most 3 in the corresponding condition were considered as potential candidates.

### Supplementary Tables

#### Table S1. iPSC clones isolated during this study

| ID | Parent cell line | Transfected construct | Mutations |
|---|---|---|---|
| WT-1 | iPS-11.10 | pSpCas9-puro + pSpCas9-hygro | - |
| WT-2 | iPS-11.10 | | - |
| WT-3 | iPS-11.10 | pSpCas9-hygro | - |
| WT-4 | iPS-11.10 | | - |
| rs11172113-141/218-4 | iPS-11.10 | pSpCas9-rs11172113-141-fw-puro + pSpCas9-rs11172113-218-rev-hygro | NC_000012.12:g.57133456_57133544del |
| rs11172113-141/218-8 | iPS-11.10 | | NC_000012.12:g.57133456_57133544del |
| rs11172113-141/218-10 | iPS-11.10 | | NC_000012.12:g.57133456_57133544del |
| rs11172113-141/218-11 | iPS-11.10 | | NC_000012.12:g.57133456_57133544del |
| rs11172113-141/218-12 | iPS-11.10 | | NC_000012.12:g.57133456_57133544del |
| LRP1KO-1 | iPS-11.10 | pSpCas9-LRP1-547-rev-puro | LRP1:c.543_563delinsA/c.545_552del |
| LRP1KO-2 | iPS-11.10 | pSpCas9-LRP1-159-rev-puro | LRP1:c.164insC/c.164insG |
| LRP1KO-3 | iPS-11.10 | | LRP1:c.164insTA/c.164del |
| LRP1KO-4 | iPS-11.10 | | LRP1:c.164insTG/c.153_165del |
| rs11172113-C/C-4 | rs11172113-141/218-10 | pSpCas9-141/218-del-puro + 1kb rs11172113-C PCR product | rs11172113-C/C |
| rs11172113-C/C-5 | rs11172113-141/218-10 | | rs11172113-C/C |
| rs11172113-T/T-7 | rs11172113-141/218-10 | pSpCas9-141/218-del-puro + 1kb rs11172113-T PCR product | rs11172113-T/T |
| rs11172113-T/T-8 | rs11172113-141/218-10 | | rs11172113-T/T |

**Table S2. Predictions of transfection binding sites affected by rs11172113 genotype.** Predictions were performed using PERFECTOS-APE webserver, using HOCOMOCO v11 (all human) transcription factor motifs database.

| Motif ID | Site rs11172113 T | P-value | Site rs11172113 C | P-value | log2 Fold Change | Gene name | Expression in arteries | Relevant publications |
|---|---|---|---|---|---|---|---|---|
| AP2B_HUMAN.H11MO.0.B | gcccAgtggc | $6.5\times10^{-4}$ | gcccGgtggc | $4.9\times10^{-5}$ | 3.7 | *TFAP2B* | - | |
| CEBPZ_HUMAN.H11MO.0.D | gcccAgtggct | $2.5\times10^{-4}$ | gcccGgtggct | $3.1\times10^{-3}$ | -3.6 | *CEBPZ* | + | |
| CXXC1_HUMAN.H11MO.0.D | cAgtggc | $1.7\times10^{-2}$ | cGgtggc | $2.5\times10^{-5}$ | 9.4 | *CXXC1* | + | |
| KLF16_HUMAN.H11MO.0.D | ttgggtgttgcccAgtggc | $1.0\times10^{-2}$ | ttgggtgttgcccGgtggc | $1.8\times10^{-4}$ | 2.5 | *KLF16* | + | |
| KLF1_HUMAN.H11MO.0.A | tgggtgttgcccAg | $3.1\times10^{-5}$ | tgggtgttgcccGg | $7.1\times10^{-6}$ | 2.1 | *KLF1* | - | |
| MECP2_HUMAN.H11MO.0.C | cccAgtg | $8.3\times10^{-3}$ | cccGgtg | $2.3\times10^{-4}$ | 5.2 | *MECP2* | + | [10,11] |
| NFYB_HUMAN.H11MO.0.A | gagccacTgggca | $4.1\times10^{-4}$ | gagccacCgggca | $5.0\times10^{-3}$ | -3.6 | *NFYB* | + | |
| NR0B1_HUMAN.H11MO.0.D | cTgggcaaca | $6.2\times10^{-3}$ | cCgggcaaca | $4.3\times10^{-4}$ | 3.8 | *NR0B1* | - | |
| RREB1_HUMAN.H11MO.0.D | attttgggtgttgcccAgtggc | $1.2\times10^{-3}$ | attttgggtgttgcccGgtggc | $2.4\times10^{-4}$ | 2.4 | *RREB1* | + | [12] |
| SNAI1_HUMAN.H11MO.0.C | cccAgtgg | $4.9\times10^{-3}$ | cccGgtgg | $3.5\times10^{-4}$ | 3.8 | *SNAI1* | + | [13] |

**Table S3. Gene ontology terms enriched for differentially expressed genes between WT and *LRP1* KO iPSC-SMCs.** ONTO: Category of gene ontology term (CC: cell component, MF: molecular function, BP: biological process), P.bonf: Bonferroni adjusted *P*-value

| ONTO | ID | Description | Gene Ratio | Background Ratio | *P*-value | *P*.bonf | *Q*-value | Included genes | Count |
|---|---|---|---|---|---|---|---|---|---|
| CC | 0062023 | collagen-containing extracellular matrix | 23/199 | 463/22283 | $3.1\times10^{-11}$ | $8.9\times10^{-9}$ | $8.5\times10^{-9}$ | *DCN/LAMC2/NTN1/FGFR2/COL16A1/ADAMTS2/LGALS1/AEBP1/CXCL12/WNT5A/TGFBI/LAMA5/FRAS1/SERPING1/ADAMTS5/LTBP3/THBS3/A2M/BCAM/EGFL6/COL15A1/COL6A6/SPON1* | 23 |
| MF | 0050840 | extracellular matrix binding | 7/202 | 57/20758 | $1.3\times10^{-6}$ | $6.1\times10^{-4}$ | $5.7\times10^{-4}$ | *DCN/LGALS1/ADGRG6/TGFBI/ITGA7/ADAMTS5/BCAM* | 7 |
| BP | 0061448 | connective tissue development | 13/196 | 288/21069 | $3.2\times10^{-6}$ | $4.4\times10^{-3}$ | $3.8\times10^{-3}$ | *SPI1/PTHLH/PPARD/HAND1/WNT5A/DLX2/TGFBI/HMGA2/FRZB/MSX1/SERPINB7/LTBP3/THBS3* | 13 |
| BP | 0030324 | lung development | 11/196 | 203/21069 | $3.3\times10^{-6}$ | $4.4\times10^{-3}$ | $3.8\times10^{-3}$ | *FGFR2/ADAMTS2/EP300/WNT5A/KLF2/LIF/LAMA5/FGF7/PDPN/ABCA3/LTBP3* | 11 |
| MF | 0005201 | extracellular matrix structural constituent | 11/202 | 199/20758 | $4.2\times10^{-6}$ | $9.9\times10^{-3}$ | $9.3\times10^{-4}$ | *DCN/LAMC2/COL16A1/AEBP1/TGFBI/LAMA5/FRAS1/THBS3/COL15A1/COL6A6/SPON1* | 11 |
| BP | 0048562 | embryonic organ morphogenesis | 13/196 | 313/21069 | $7.9\times10^{-6}$ | $6.5\times10^{-3}$ | $5.7\times10^{-3}$ | *NTN1/FGFR2/EYA1/HAND1/WNT5A/DLX2/TBX3/CRB2/FRZB/MSX1/RBPMS2/IRX5/GATA2* | 13 |
| BP | 0048332 | mesoderm morphogenesis | 7/196 | 87/21069 | $1.7\times10^{-5}$ | $9.2\times10^{-3}$ | $8.0\times10^{-3}$ | *FGFR2/EYA1/HAND1/WNT5A/TBX3/CRB2/HMGA2* | 7 |
| CC | 0005604 | basement membrane | 7/199 | 99/22283 | $3.0\times10^{-5}$ | $4.4\times10^{-3}$ | $4.2\times10^{-3}$ | *LAMC2/NTN1/TGFBI/LAMA5/FRAS1/EGFL6/COL15A1* | 7 |
| MF | 0005539 | glycosaminoglycan binding | 11/202 | 262/20758 | $5.5\times10^{-5}$ | $8.5\times10^{-3}$ | $8.0\times10^{-3}$ | *DCN/LAMC2/FGFR2/CEMIP/PTPRS/CCL7/FGFBP1/FGF7/ADAMTS5/THBS3/GREM2* | 11 |

#### Supplementary Figures

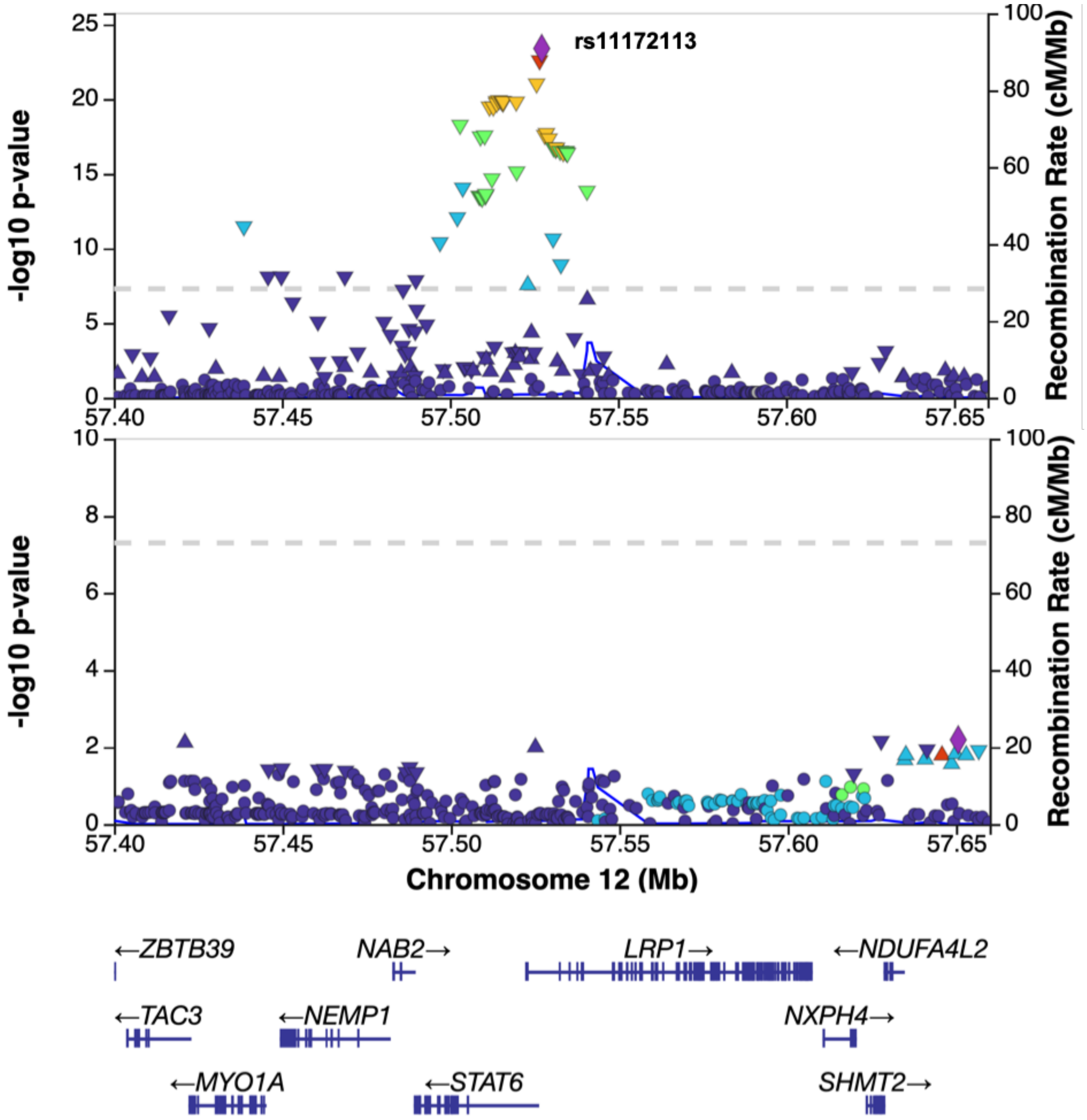


**Figure S1. Conditional association analysis on rs11172113 with migraine at *LRP1* locus.** Representative locusZoom plots of migraine association signal before (top) and after (bottom) conditioning on the association with rs11172113 using GCTA COJO method[26].

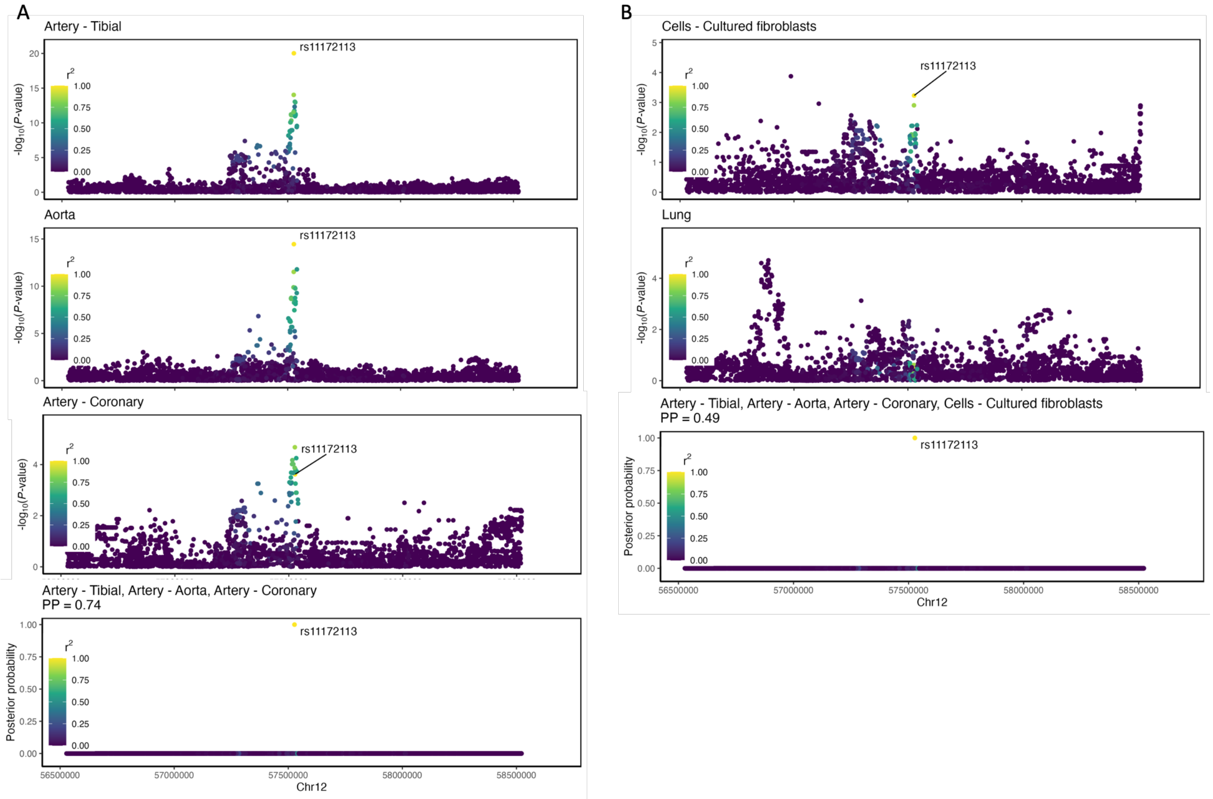


**Figure S2. eQTL colocalization at LRP1 locus. A-B:** *LRP1* eQTL signals in Tibial artery, Aorta, Coronary artery (A) Cultured fibroblasts and Lung (B) are represented in a 2Mb region centered on rs11172113. Association P-value (log10 scale) of each SNP is represented on y-axis, while dot color represents $r^2$ of linkage disequilibrium with rs11172113 (European population of 1000G reference panel). Lower panels represent the relative posterior probability for each SNP to be causal, while the total posterior probability for the associations to colocalize at the locus is given over the graph.

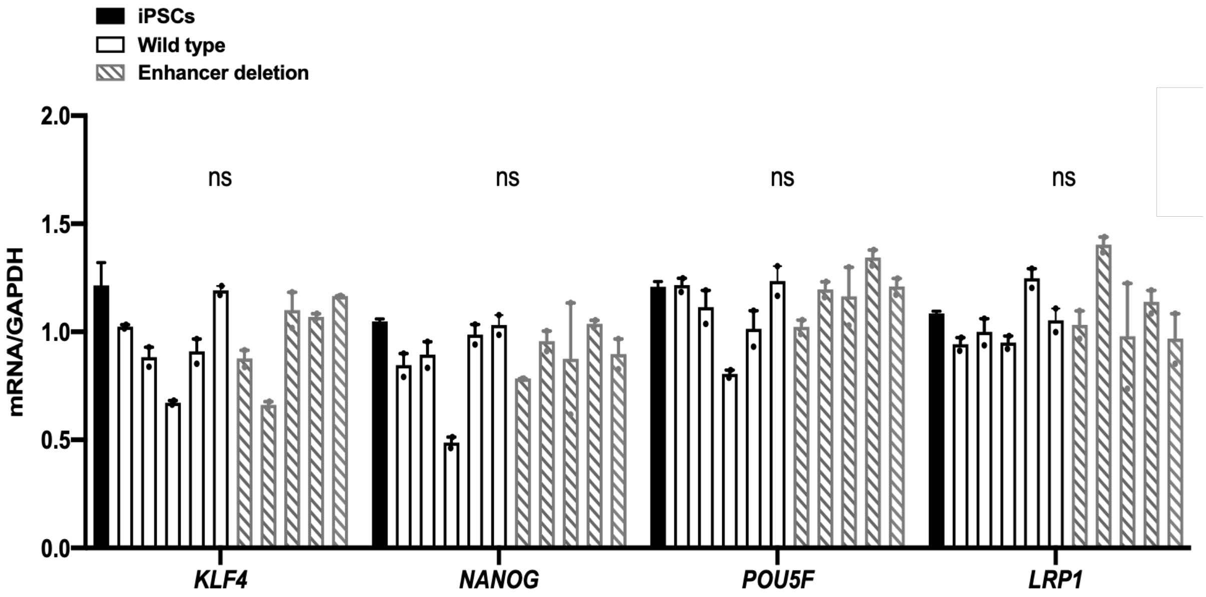


**Figure S3. Pluripotent markers in iPSCs with or without rs11172113 enhancer deletion.** Bar plots represent mean±SEM of mRNA expressions of pluripotent marker genes and *LRP1* in unmodified, wild types, enhancer deleted iPSCs (grey with stripes). Unadjusted t-test: ns: not significant.

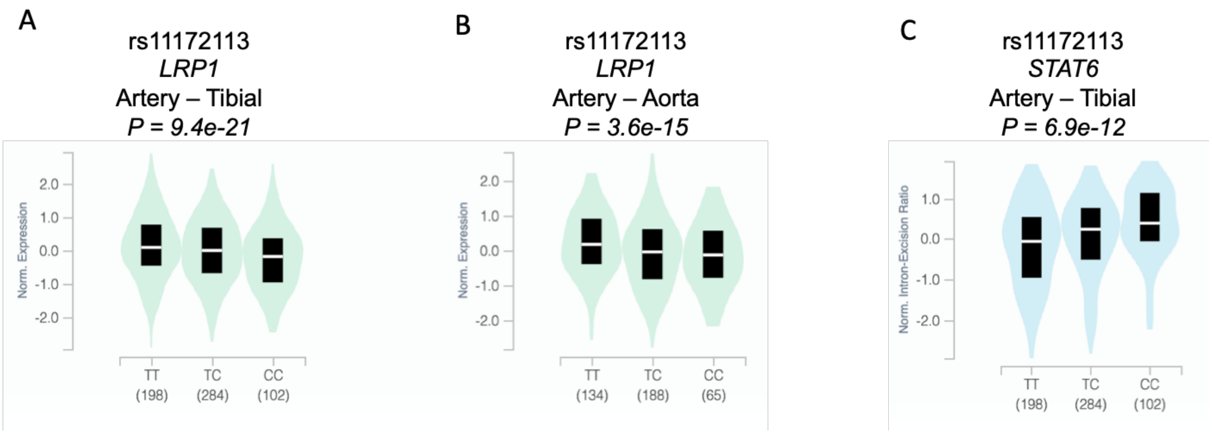


**Figure S4. Genotype-expression correlation of *LRP1* and *STAT6* with rs11172113. A-B:** Violin plots showing normalized expression of *LRP1* in tibial artery (**A)** and aorta **(B**) depending on rs11172113 genotype in GTEx data. **C**: Relative detection of *STAT6* exon junction 57108299:57111129 (exon1-exon2) depending on rs11172113 genotype in tibial artery tissue.

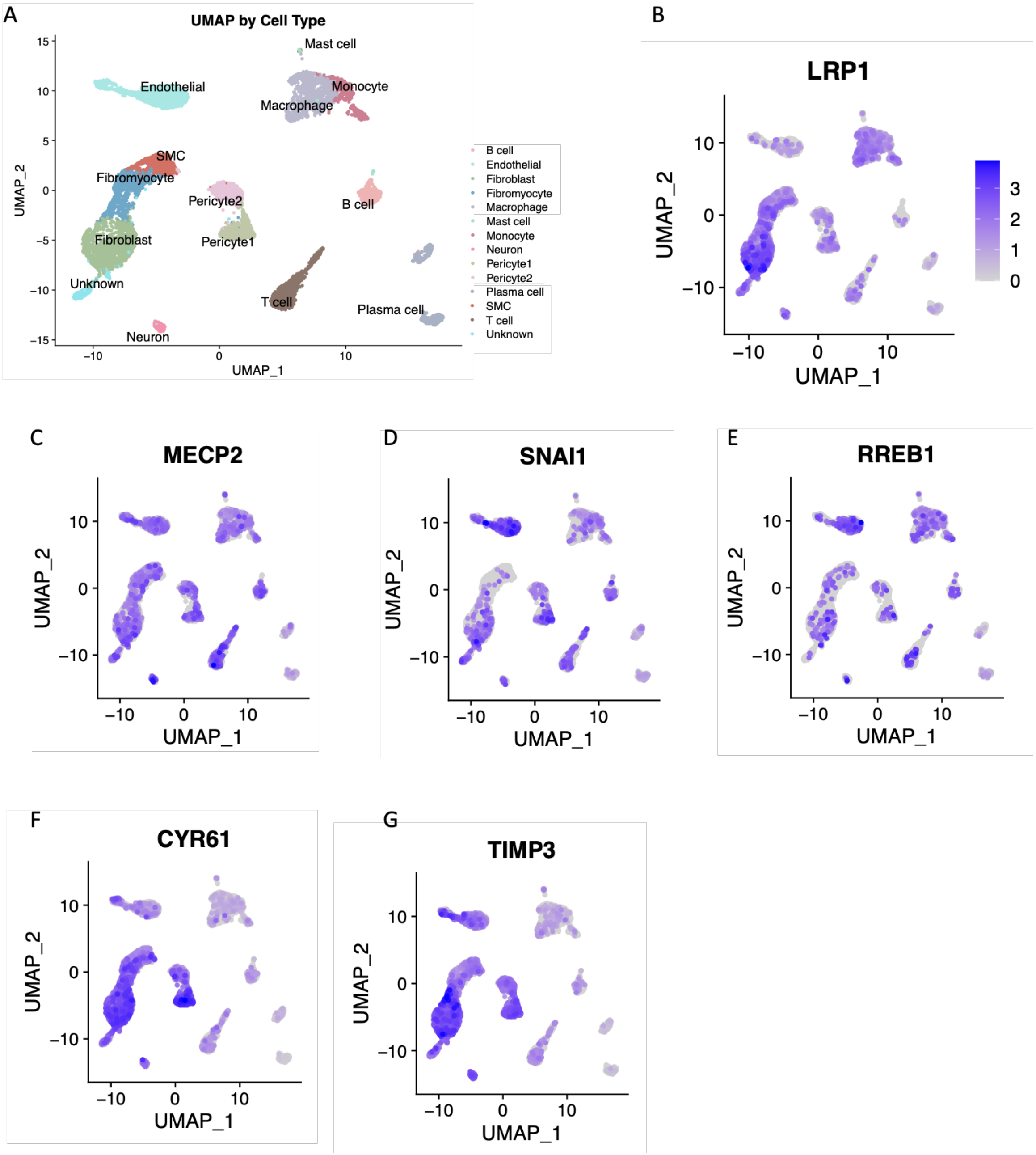


**Figure S5. Single cell expression in human coronary arteries**

**A**: Representative UMAP plots showing single cell populations profiled in single nuclei RNA-Seq analysis of diseased human coronary arteries[14] visualized using PlaqView.[15] **B-G** Featureplots of *LRP1* (**B**), *MECP2* (**C**), *SNAI1* (**D**), *RREB1* (**E**), *CYR61* (**F**), *TIMP3* (**G**).

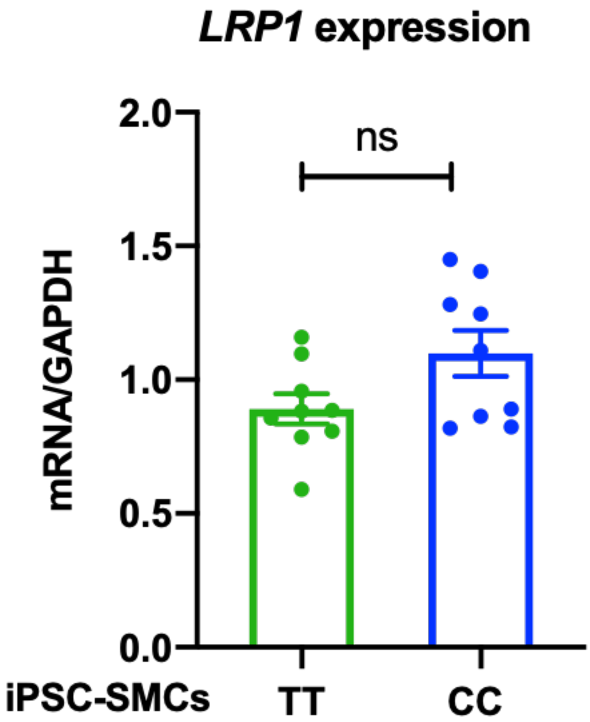


**Figure S6. LRP1 expression in iPSC-SMCs homozygous for rs11172113.** RNA levels of *LRP1* detected in iPSC-SMCs with TT (green) or CC genotypes (blue) generated using CRISPR/Cas9. 2 clones for each genotype were analyzed with four replicates. Unadjusted t-test: ns: not significant.

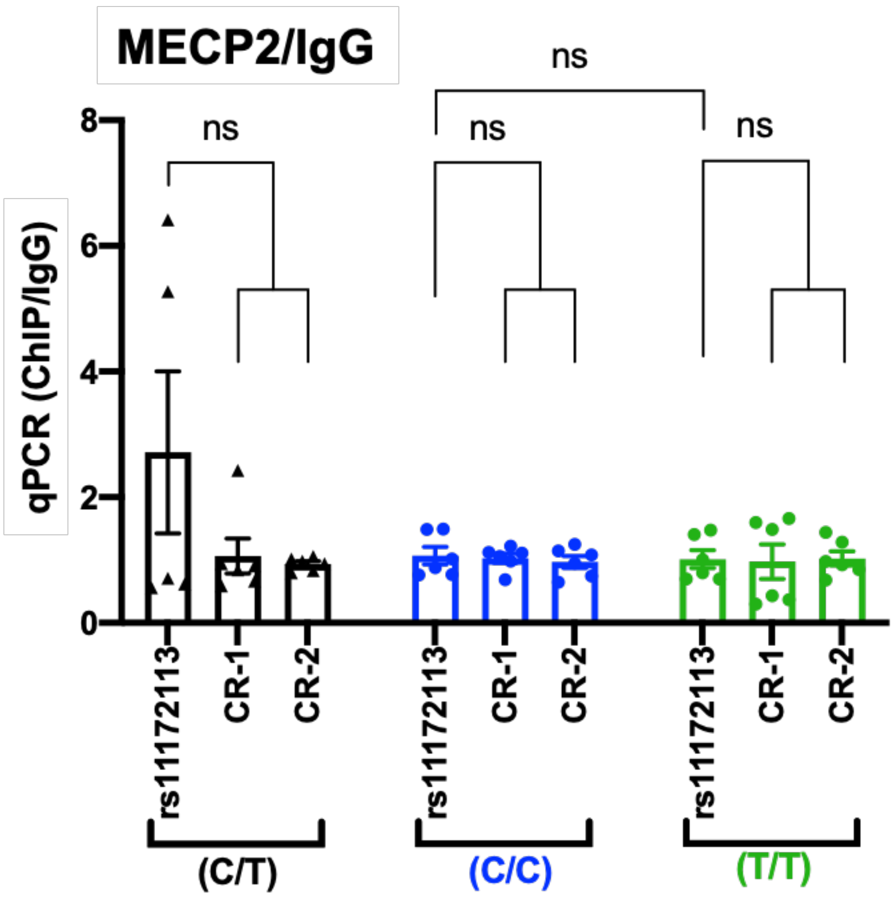


**Figure S7. MECP2 binding to rs11172113 region.** Chromatin immunoprecipitation (ChIP) targeting MECP2 in iPSC-11.10 and homozygous rs11172113 C/C or T/T (two clones each). Immunoprecipitated material was evaluated using qPCR targeting and rs11172113 region and two control regions (no enhancer marks in SMCs or artery tissue). The ratio of MECP2 ChIP to ChIP with rabbit IgG is given. Each clone was assessed in triplicate. Unadjusted T-test: *p<0.05, **p<0.01, ns: not significant.

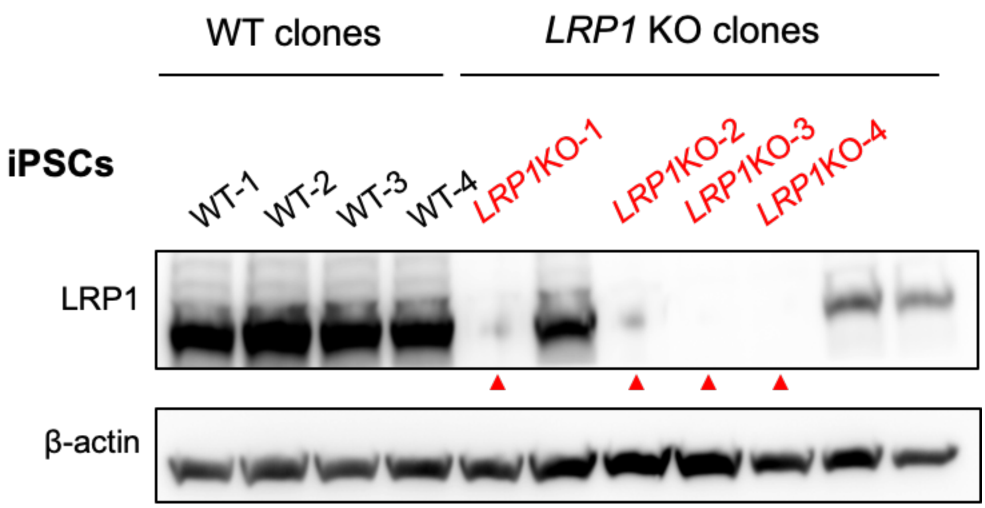


**Figure S8. LRP1 protein expression in *WT* and *LRP1*-KO iPSCs.** Representative images of Western blots of LRP1 expression in WT and screened clones transfected with sgRNA targeting LRP (“LRP1 KO clones”). Clones with detected LRP1 expression were discarded and are not named. Genotype of clones with no LRP1 expression was confirmed using Sanger sequencing (Table S1)

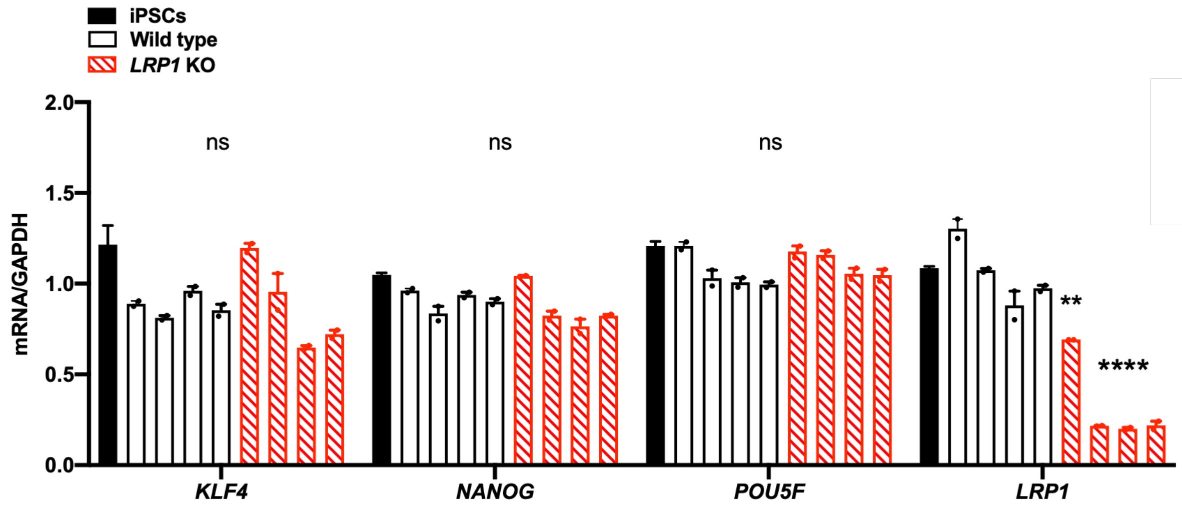


**Figure S9. Expression of pluripotent cells markers in WT and *LRP1* KO clones.** Bar plots represent mean±SEM of relative mRNA expressions of pluripotent marker genes and *LRP1* in unmodified, wild types, and *LRP1* KO (red with stripes) iPSCs. Unadjusted t-test comparing each clone to parent iPSC cell line:, **$p<0.05$, ****$p<10^{-4}$, ns: not significant.

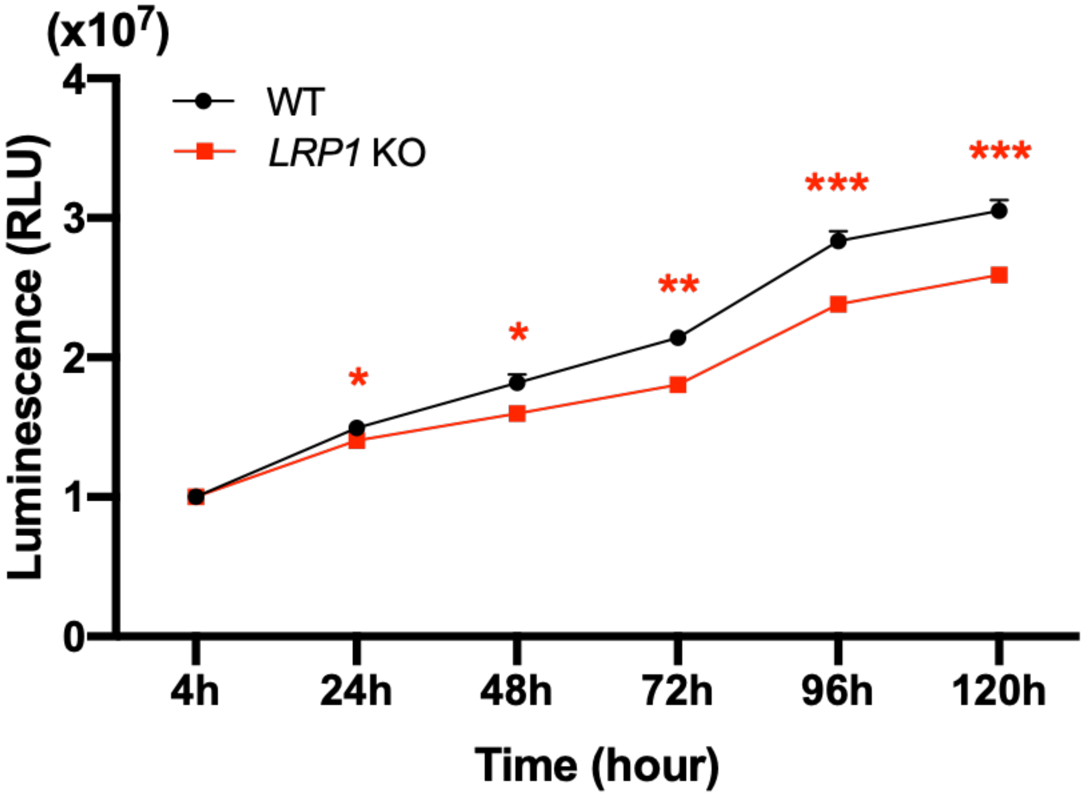


**Figure S10. Cell proliferation in WT and LRP1 KO iPSC-SMCs.** Measurement of cell viability(mean±SEM,) of WT (black dots, 4 clones) and *LRP1* KO SMCs (red squares, 3 clones) at 4h, 24h, 48h, 72h, 96h and 120h. The unadjusted t-test was used at each time point: *$p<0.05$, **$p<0.01$, ***$p<0.001$, ns: not significant.

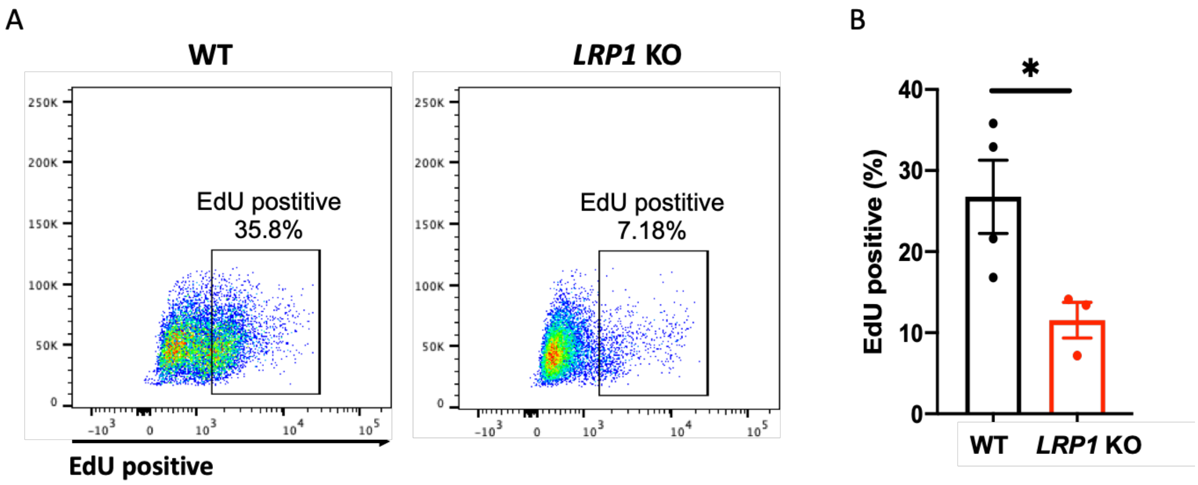


**Figure S11. DNA replication in WT and LRP1 KO iPSC-SMCs. A**: Representative plots of fluorescence-activated cell sorting (FACS) exhibiting EdU incorporation into cells. **B**: Bar plots showing the quantification of EdU positive cells (mean±SEM, 4 WT and 3 KO clones). Unadjusted t-test: *p<0.05.

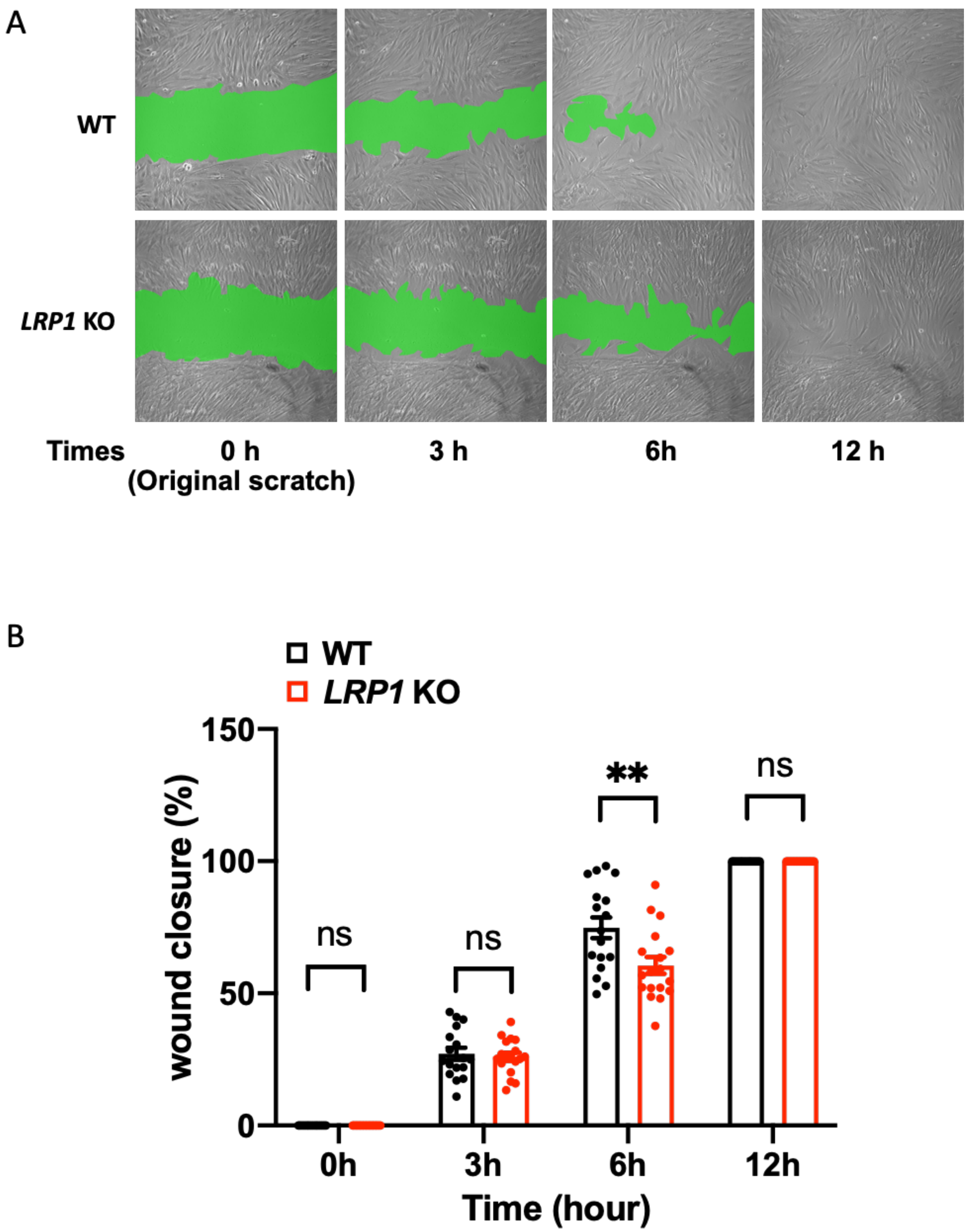


**Figure S12. Wound closure in WT and LRP1 KO iPSC-SMCs. A**: Representative images from scratch wound repair assay in of WT and *LRP1* KO SMCs at 0h (original scratch), 3h, 6h and 12h. Green area represents the wound area used for measurements. **B**: Barplot representing mean±SEM of the percentage of wound closure at 3h, 6h and 12h (N=16).

Wound area (green area) was measured by Image J and compared to wound area measured at 0h. Unadjusted t-test:**$p<0.01$, ns: not significant.

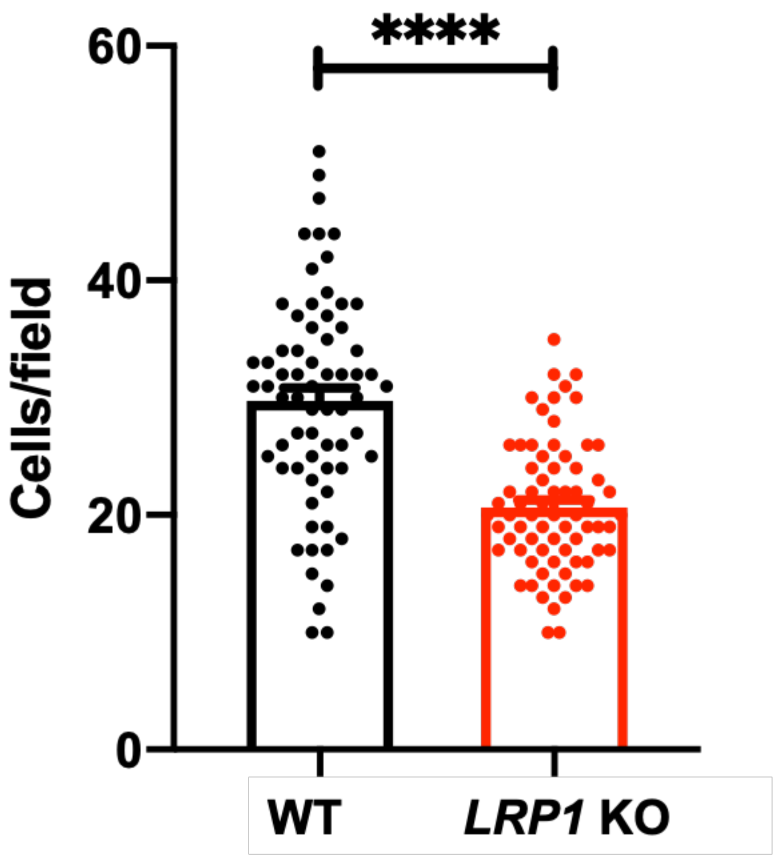


**Figure S13. Transwell cell migration assay in WT and *LRP1* KO iPSC-SMCs.** Barplot representing mean±SEM of the number of cells counted in each field following Transwell assay assessing cell migration through polycarbonate membrane (8µm pore size, 20h culture, 67 fields from 6 independent replicates). Unadjusted t-test: ****p<0.0001.

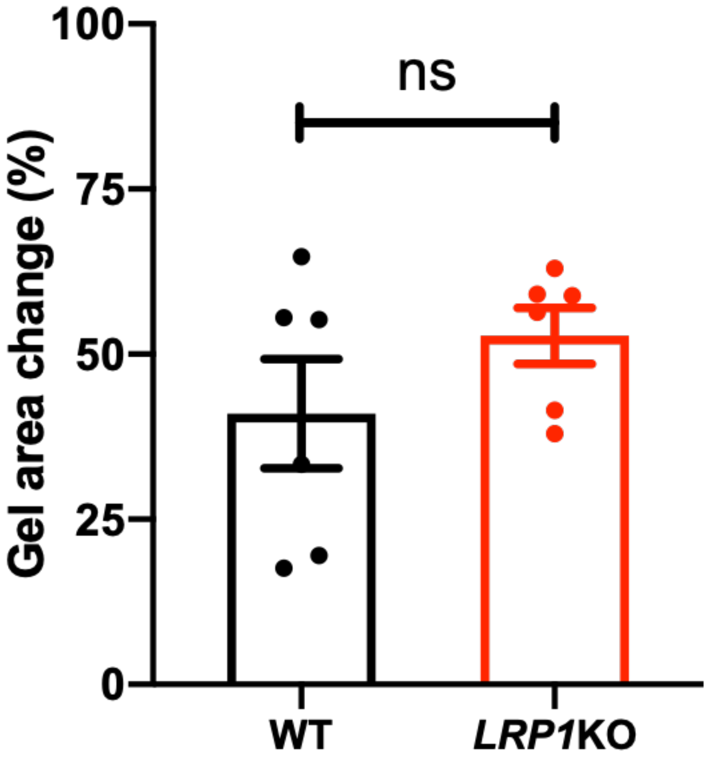


**Figure S14. Collagen gel contraction by WT and *LRP1* KO iPSC-SMCs.** Representative barplot represent the mean±SEM of reduction in gel area of collagen gel lattices with no cells or WT and *LRP1* KO SMCs after 20 h (N=6). Unadjusted t-test: ns: not significant.

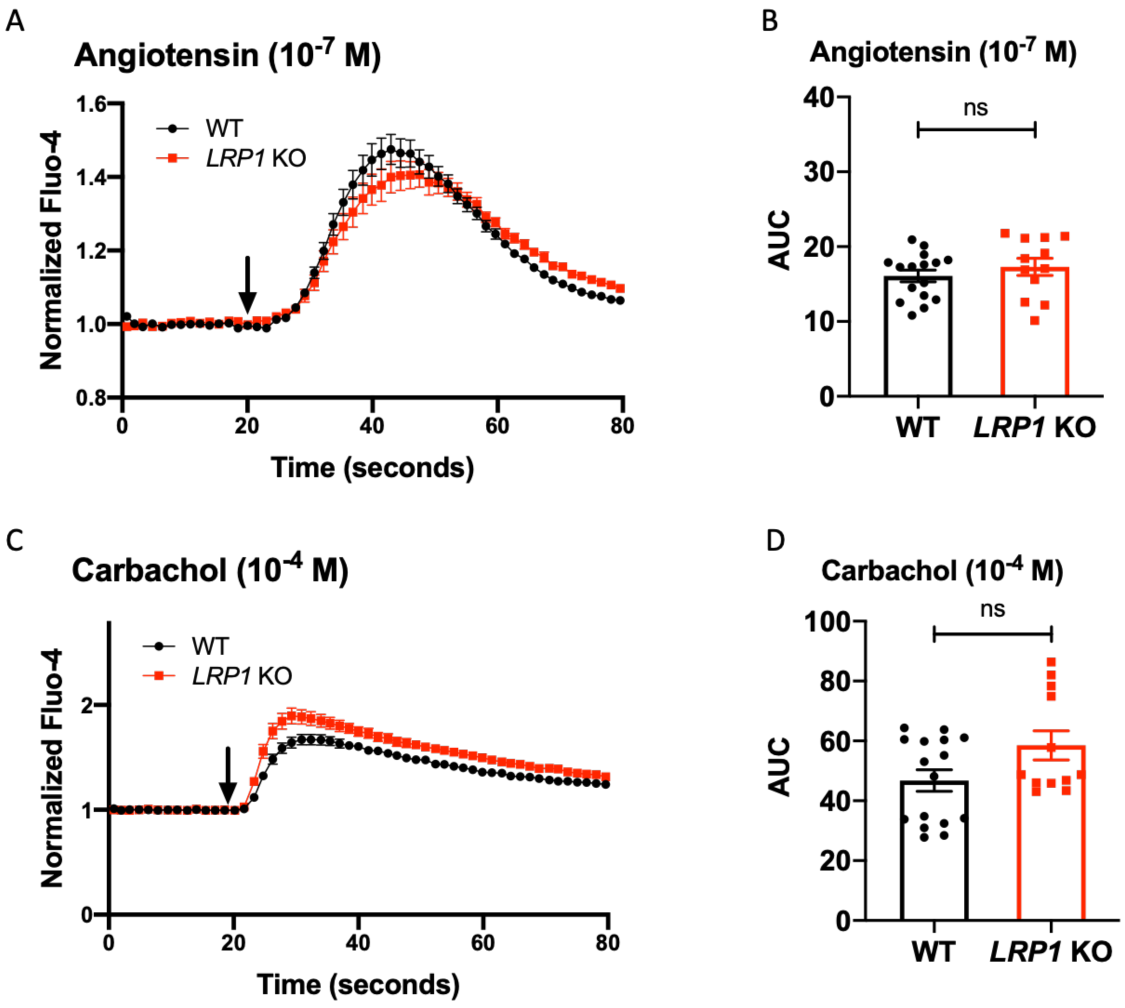


**Figure S15. Calcium release in WT and *LRP1* KO iPSC-SMCs.** Measure of fluorescence in cells labeled with intracellular calcium probe fluo-4 and treated by Angiotensin II (**A**) or Carbachol (**C**) as indicated in WT (black) and *LRP1* KO SMCs (red). The arrow represents the time point at which the inducer was added to the cells. Representative graph showing calculated area under the curve (AUC) for Angiotensin II (**B**) or Carbachol (**D**). Comparison of AUC were performed using unadjusted t-test as indicated: n=16, mean±SEM, ns: not significant.

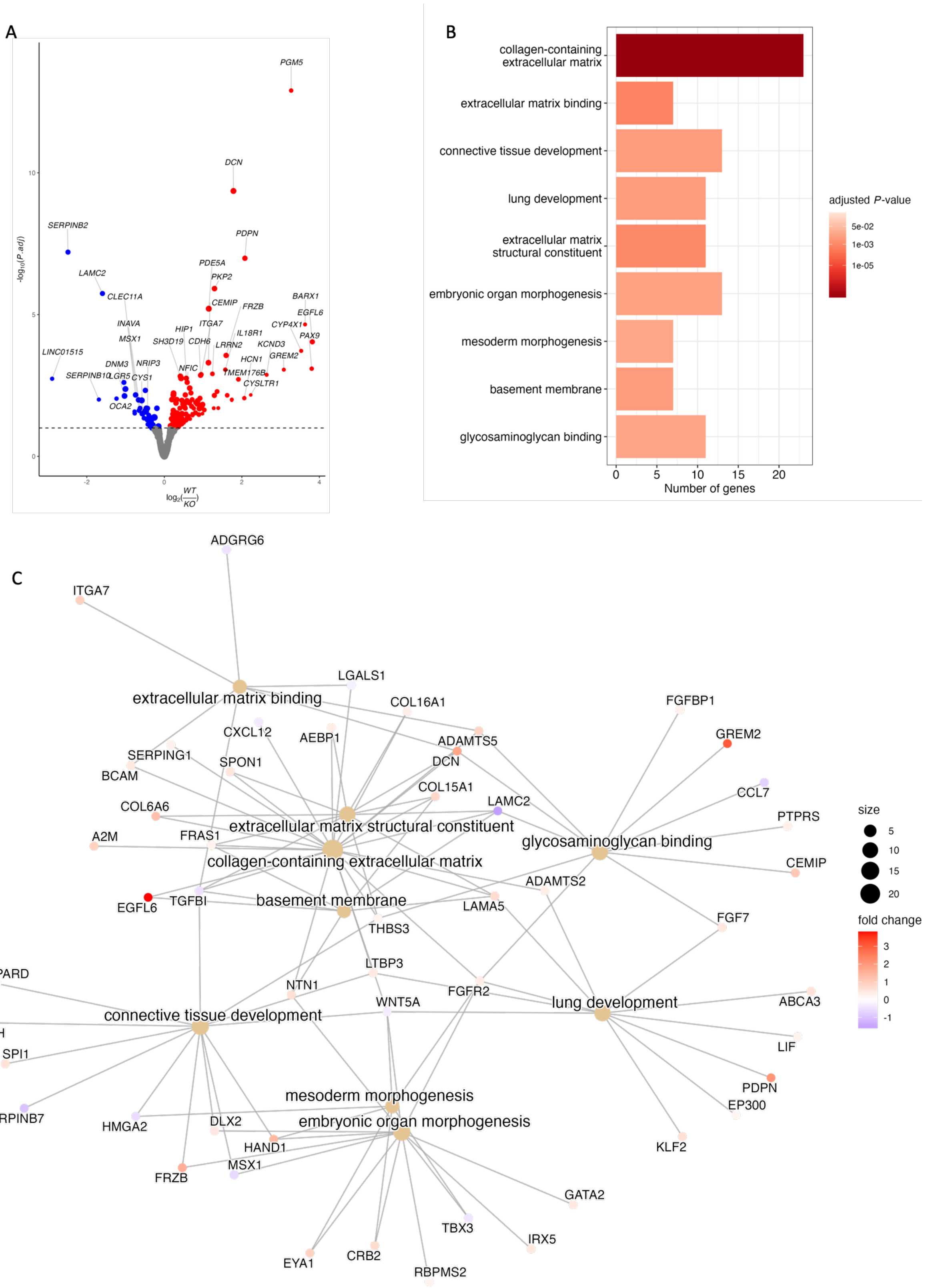


**Figure S16. RNA-Seq analysis of differential gene expression in WT and *LRP1* KO iPSC-SMCs. A:** Volcano plot representing P-value of differential expression (y-axis, log10 scale) versus estimated log2 Fold Change (WT over LRP1 KO). Genes significantly

(FDR<0.1) overexpressed in WT iPSC-SMCs are represented in red, while genes overexpressed in *LRP1* KO cells are in blue. Name of top differentially expressed genes is indicated. **B:** Barplot representing number of genes and adjusted P-valu of top enriched Gene Ontology (GO) termes involving differentially expressed geneS. **C:** Cnet plot of GO terms enriched in differentially expressed genes.

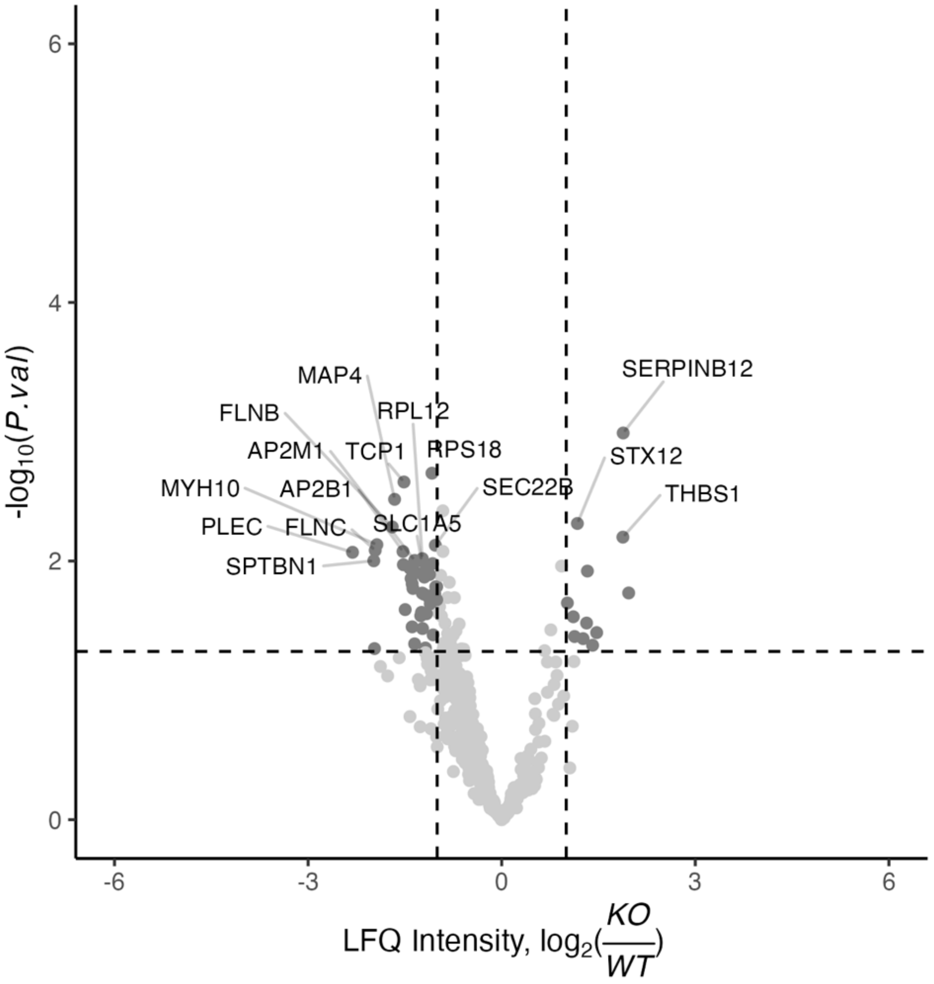


**Figure S17. Quantitative assessment of extracellular matrix generated by WT and *LRP1* KO iPSC-SMCs.** Volcano plot representing P-value of differential protein expression (y-axis, log10 scale) versus estimated log2 Fold Change (*LRP1* KO over WT). Nominally significant differentially expressed proteins (*P*<0.05 and |log2 Fold Change| > 1) overexpressed in WT iPSC-SMCs are represented in dark grey, but no protein was statistically significant after adjusting for multiple testing. Name of top differentially expressed proteins is indicated.

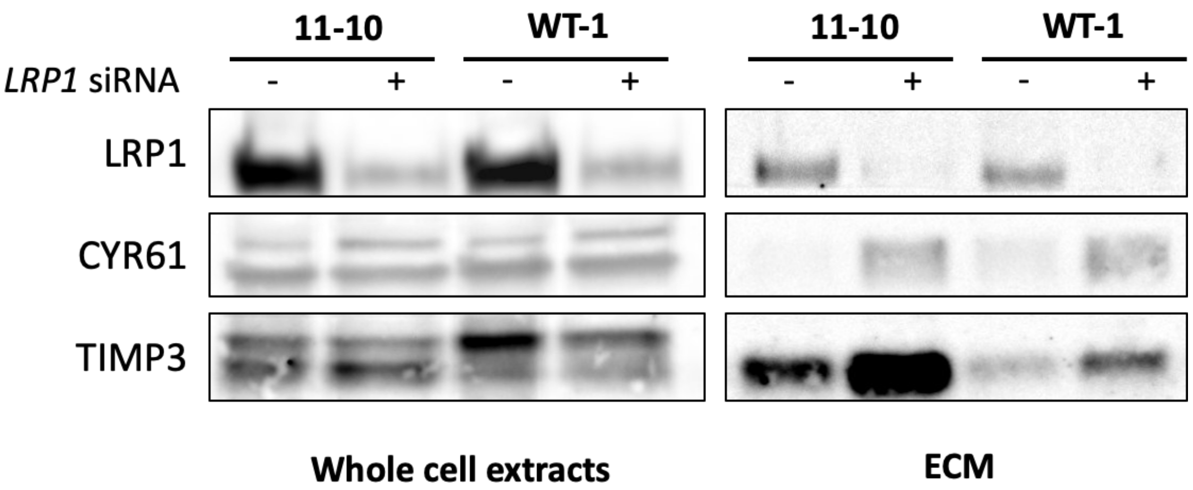


**Figure S18. Extracellular matrix (ECM) protein expression following *LRP1* knockdown in iPSC-SMCs.** Representative images of Western blots of LRP1, CYR61 and TIMP3 expression in whole cell extracts or ECM only in the presence of siRNA targeting *LRP1*.
